# Uncontrolled CD21^low^ age-associated and B1 B cell accumulation caused by failure of an EGR2/3 tolerance checkpoint

**DOI:** 10.1101/2021.07.01.450650

**Authors:** Etienne Masle-Farquhar, Timothy J. Peters, Lisa A. Miosge, Ian A. Parish, Christoph Weigel, Christopher C. Oakes, Joanne H. Reed, Christopher C. Goodnow

**Author notes:** Equal senior authors (Joanne H. Reed, Christopher C. Goodnow).

## Abstract

CD21^low^ age-associated or atypical memory B cells, enriched for autoantibodies and poised for plasma cell differentiation, accumulate in large numbers in chronic infections, autoimmune disease and immunodeficiency, posing the question of what checkpoints normally oppose their excessive accumulation. Here, we reveal a critical role for the calcium-NFAT-regulated transcription factors EGR2 and EGR3. In the absence of EGR2 and EGR3 within B cells, CD21^low^ and B1 B cells accumulate and circulate in young mice in numbers 10-20 times greater than normal, over-express a large set of EGR2 ChIP-seq target genes including known drivers of plasma cell differentiation and under-express drivers of follicular germinal centers. Most follicular B cells constitutively express *Egr2* proportionally to surface IgM down-regulation by self-antigens, and EGR2/3 deficiency abolishes this characteristic anergy response. These results define a key transcriptional checkpoint repressing CD21^low^ B cell formation and inform how *NFATC1* or *EGR2* mutations promote B1 cell-derived chronic lymphocytic leukemias.

## INTRODUCTION

Atypical CD21^low^ B cell populations accumulate in large numbers in multiple disease states, but little is known about the physiological checkpoints normally opposing their formation. These cells, variously labelled atypical-memory, IgD/CD27 double-negative, age-associated or CD21^low^ B cells, share the unusual loss of cell-surface CR2 complement C3d receptor (CD21) expression, and display gene expression and antibody V-region profiles intermediate between follicular or memory B cells on the one hand, and plasmablasts on the other (Charles et al., 2011, Isnardi et al., 2010, Jenks et al., 2018, Rakhmanov et al., 2009, Rubtsov et al., 2011, Russell Knode et al., 2017, Terrier et al., 2011, Scharer et al., 2019). We refer to them collectively here as CD21^low^ B cells.

Atypical CD21^low^ B cells were first found accumulating in individuals with human immunodeficiency virus (HIV) chronic viremia (Benedetto et al., 1992, Moir et al., 2001), and contain the majority of cells bearing immunoglobulin against HIV envelope gp120 (Moir et al., 2008). CD21^low^ B cells also accumulate in individuals with common variable immunodeficiency (CVID) and concomitant autoimmune cytopenias (Isnardi et al., 2010, Warnatz et al., 2002) or severe systemic lupus erythematosus (SLE) (Rakhmanov et al., 2009, Wei et al., 2007). CD21^low^ B cells that accumulate with chronic HCV infections often express rheumatoid factor IgM that binds self-IgG (Charles et al., 2011, Terrier et al., 2011) and in CVID and lupus, many CD21^low^ B cells express self-reactive *IGHV4-34* immunoglobulins (Rakhmanov et al., 2009, Wei et al., 2007). CD21^low^ CD23^low^ B cells, many expressing anti-chromatin immunoglobulins (Russell Knode et al., 2017), accumulate in the archetypal systemic autoimmune mouse strain, NZB/W, in *Mertk*^*-/-*^mice genetically predisposed to lupus-like autoimmunity, or in old C57BL/6 mice without overt autoimmune disease (Hao et al., 2011, Rubtsov et al., 2011).

Here, we reveal the critical role in opposing CD21^low^ B cell accumulation served by Early Growth Response 2 (EGR2) and EGR3. These paralogous zinc finger transcription factors are encoded by immediate early genes that are induced by chronic self-antigen stimulation of cell-surface IgM and IgD through calcium- and calcineurin-activated transcription factors of the nuclear factor of activated T cells (NFAT) family (Healy et al., 1997, Glynne et al., 2000, Merrell et al., 2006, Sabouri et al., 2016, Marklin et al., 2017). The function of EGR2 and EGR3 in B cells is not known, as earlier knockout mouse studies were confounded by their essential roles in T cells (Li et al., 2012). *EGR2* is induced in CD21^low^ B cells in humans (Isnardi et al., 2010, Terrier et al., 2011) and mice (Russell Knode et al., 2017), and somatic missense *EGR2* mutations recur and predict poor outcome in B cell chronic lymphocytic leukemia (CLL)(Young et al., 2017). CLLs display constitutive NFAT activation(Schuh et al., 1996), resemble self-reactive anergic B cells, and derive from another chronically-stimulated self-reactive population – B1 cells(Apollonio et al., 2013). Like CD21^low^ B cells, B1 cells also accumulate in large numbers in NZBW autoimmunity-prone mice (Hayakawa et al., 1983). *EGR2* is also induced by *acute* BCR stimulation to promote B cell proliferation *in vitro* (Glynne et al., 2000, Li et al., 2012, Newton et al., 1996), highlighting the question of what functions EGR2 and EGR3 perform in B cells *in vivo*.

Here, we studied the functions of EGR2/3 in B cells *in vivo* by analysing chimeric mice with *Egr2* and *Egr3* deletion restricted to a fraction of B cells. This revealed a critical, cell-autonomous function for EGR2/EGR3 in repressing formation of CD21^low^ and B1 B cell populations, and in promoting the IgM^low^ IgD^high^ anergy response of self-reactive follicular B cells. Using flow cytometry and single-cell RNA sequencing, we define the landscape of EGR2/EGR3-repressed and -induced genes in CD21^low^, B1a and follicular B cells, many corresponding to EGR2 chromatin immunoprecipitation sequencing (ChIP-seq) targets in CLL. Our findings define a critical EGR2/EGR3 transcriptional checkpoint governing accumulation of CD21^low^ and B1 B cells and reveal the molecular circuit responsible for the cardinal features of anergic B cells.

## RESULTS

### Egr2 and Egr3 repress mature CD21^low^ CD23^low^ B cells

To study the functions of *Egr2* and *Egr3* in B cells, we intercrossed germline *Egr3* knockout mice with mice expressing a loxP-flanked *Egr2* allele and a *Cre* transgene controlled by the *Cd19* promoter, resulting in B cell-specific *Egr2* deletion. All mice examined were matched for *Cd19*^*Cre*/+^ and we thus describe *Egr2*^*fl*^ *Egr3*^*KO*^ mice based only on their *Egr2* and *Egr3* genotypes.

Mice lacking one or both alleles of *Egr2* and/or *Egr3* in B cells had no changes in the bone marrow with respect to mean frequency or number of precursor B cells, immature B cells or mature recirculating B cells (**Supplementary Figure 1A**,**B**). In the spleen, *Egr2*^*fl/fl*^ *Egr3*^*KO/KO*^ mice had a 3-fold increase in mean number of CD93^pos^ CD23^neg^ immature T1 B cells (**Supplementary Figure 1C**), many of them with low cell-surface IgM expression (**Supplementary Figure 1D**).

The most striking abnormality in the spleen of *Egr2*^*fl/fl*^ *Egr3*^*KO/KO*^ mice was a 30- and 10-fold increase in mean frequency and number of B220^pos^ CD19^pos^ CD95^neg^ CD93^neg^ mature B cells with a CD21^low^ CD23^low^ phenotype (**Figure 1A**, **B**). Other B cell (CD19^pos^) subsets that can express low levels of CD21 and CD23 were excluded from this analysis: B1a cells (B220^int^ CD5^high^ CD43^high^ CD23^low^), germinal centre B cells (IgD^neg^ CD38^low^ CD95^pos^) and immature B cells (CD93^pos^). CD21^low^ CD23^low^ mature B cells were also expanded a mean 16-fold in the blood (**Figure 1C**) and 5-fold in the bone marrow (**Supplementary Figure 2A**) of *Egr2*^*fl/fl*^ *Egr3*^*KO/KO*^ relative to wild-type mice. A subtle increase in CD21^low^ CD23^low^ mature B cells occurred in *Egr2*^*fl/fl*^ *Egr3*^*KO/+*^ mice retaining one functional *Egr3* allele, but homozygous loss of *Egr2* or *Egr3* alone had no discernible effect (**Figure 1B**,**C; Supplementary Figure 2A**). The mean numbers of mature follicular and marginal zone B cells were normal in *Egr2*^*fl/fl*^ *Egr3*^*KO/KO*^ mice (**Figure 1B**), but mean cell-surface CD23 and CD21 expression were decreased on mature recirculating B cells in the bone marrow, and on immature T2 and T3 cells and splenic follicular B cells in the spleen of *Egr2*^*fl/fl*^ *Egr3*^*KO/KO*^ mice (**Supplementary Figure 2B**).

**Figure 1.**
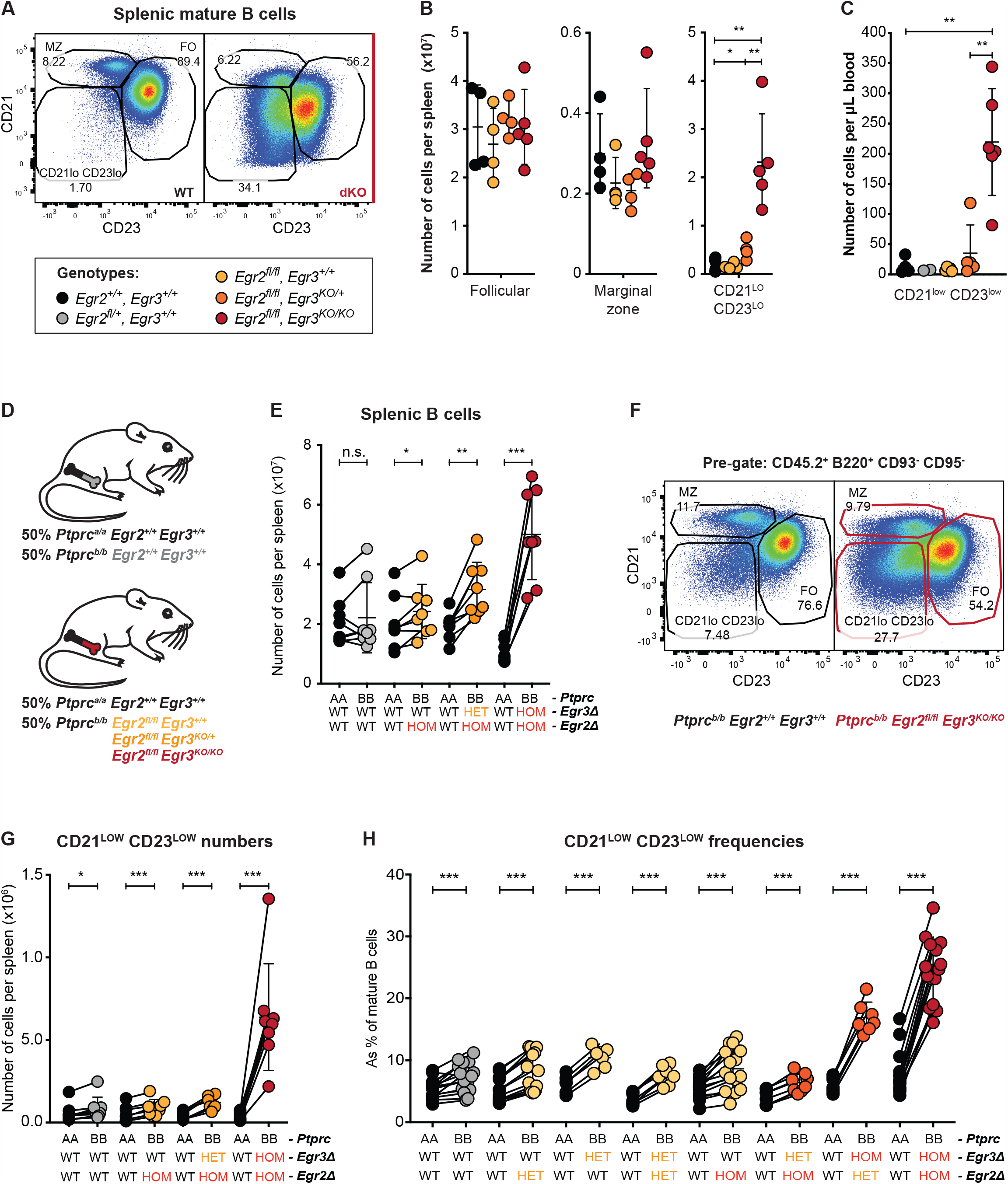
Cell-autonomous accumulation of *Egr2* and *Egr3* deficient CD21^low^ CD23^low^ age-associated-like B cells. **A**, Flow cytometric analysis of splenic B220^pos^ CD95^neg^ CD93^neg^ mature CD23^pos^ follicular (FO), CD23^low^ CD21^pos^ marginal zone (MZ) and CD21^low^ CD23^low^ B cells, in *Egr2*^*+/+*^ *Egr3*^*+/+*^ (WT) or *Egr2*^*fl/fl*^ *Egr3*^*KO/KO*^ (dKO) mice. **B**, Total number per spleen of follicular, marginal zone or CD21^low^ CD23^low^ B cells in *Egr2*^*+/+*^ *Egr3*^*+/+*^ (black), *Egr2*^*fl/fl*^ *Egr3*^*+/+*^ (yellow), *Egr2*^*fl/fl*^ *Egr3*^*KO/+*^ (orange) and *Egr2*^*fl/fl*^ *Egr3*^*KO/KO*^ (red) mice. **C**, Number of CD21^low^ CD23^low^ B cells per μL blood, in mice of the indicated genotypes. **D**, *Ptprc, Egr2* and *Egr3* genotypes of bone marrow transplanted into *Rag1*^*KO/KO*^ mixed to generate chimeric mice. **(E, G, H)** Each circle represents one mouse. Black lines denote cells within one chimeric mouse. **E**, Total number per spleen of *Ptprc*^*a/a*^ or *Ptprc*^*b/b*^ B cells, in mixed chimeric mice. **F**, Flow cytometric analysis of CD45.2^pos^ FO, MZ and CD21^low^ CD23^low^ B cells, in mice that received *Egr2*^*+/+*^ *Egr3*^*+/+*^ (left) or *Egr2*^*fl/fl*^ *Egr3*^*KO/KO*^ (right) *Ptprc*^*b/b*^ bone marrow. **G**, Total number per spleen of *Ptprc*^*a/a*^ or *Ptprc*^*b/b*^ CD21^low^ CD23^low^ B cells, in mixed chimeric mice. **H**, Percentage of *Ptprc*^*a/a*^ or *Ptprc*^*b/b*^ mature B cells with a CD21^low^ CD23^low^ phenotype, in mixed chimeric mice. **(A-H)** Data are represented as mean ± SD. **(A-C)** Data representative of *n* = 3 experiments in mice 8-20 weeks old. Comparisons made by multiple *t*-tests, with Holm-Šidák correction. **(D-H)** Data pooled from *n* > 2 independent experiments with *n* = 4 recipients per group. Comparisons made by paired t-test. * p < 0.05; ** p < 0.01; *** p < 0.001.

The accumulation of CD21^low^ CD23^low^ cells in *Egr2*^*fl/fl*^ *Egr3*^*KO/KO*^ mice could reflect indirect effects of *Egr3* deficiency in non-hematopoietic cells. To address this, we generated “100% chimeras” by irradiating *Rag1*^*KO/KO*^ mice and transplanting bone marrow from an *Egr2*^*fl/fl*^ *Egr3*^*KO/KO*^ mouse or from *Egr2*^*+/+*^ *Egr3*^*+/+*^ or *Egr2*^*fl/+*^ *Egr3*^*KO/+*^ control donors. *Egr2*^*fl/fl*^ *Egr3*^*KO/KO*^ marrow recipients had no significant change in frequency of immature or mature B cells (**Supplementary Figure 3A**), but had an 8-fold and 10-fold increase in mean frequency of CD21^low^ CD23^low^ B cells in the spleen and bone marrow, respectively (**Supplementary Figure 3B**). Homozygous *Egr2* and *Egr3* deficiency restricted to B cells and hematopoietic cells is therefore sufficient to cause accumulation of CD21^low^ CD23^low^ B cells.

To determine the *B cell*-intrinsic effects of *Egr2* and *Egr3* deficiency, we generated mixed chimeras by irradiating *Rag1*^*KO/KO*^ mice and transplanting an equal mixture of “test” bone marrow from *Ptprc*^*b/b*^ *Cd19*^*Cre/+*^ mice lacking one or both alleles of *Egr2* and *Egr3* and control bone marrow from *Ptprc*^*a/a*^ *Cd19*^*+/+*^ *Egr2*^*+/+*^ *Egr3*^*+/+*^ donors (lower panel in **Figure 1D**). As an additional control, some mixed chimeras received *Ptprc*^*b/b*^ *Cd19*^*Cre/+*^ “test” marrow with wild-type *Egr2* and *Egr3* genes (upper panel in **Figure 1D**). We stained cells in the reconstituted chimeras with antibodies to CD45.1 or CD45.2 (encoded by *Ptprc*^*a*^ or *Ptprc*^*b*^, respectively) and applied an identical gating strategy for CD45.1^pos^ control leukocytes and CD45.2^pos^ test leukocytes. CD45.2+ B cells with homozygous *Egr2* and *Egr3* deletion accumulated in the spleen in greater numbers than CD45.1+ wild-type control B cells in the same animal (**Figure 1E**). In addition, we observed a 20-fold accumulation of CD21^low^ CD23^low^ B cells restricted to cells lacking EGR2 and EGR3 (**Figure 1F, G**), establishing that these transcription factors act cell-autonomously to inhibit this atypical B cell population.

To determine the relative effects of *Egr2* versus *Egr3* deletion on CD21^low^ CD23^low^ B cell accumulation, we combined data from a series of mixed chimera experiments. Heterozygous *Egr2* or *Egr3* deletion alone or combined, and homozygous *Egr2* deficiency coupled with heterozygous *Egr3* deletion, were insufficient to expand CD21^low^ CD23^low^ B cells when compared to the additional control of *Ptprc*^*b/b*^ *Cd19*^*Cre/+*^ *Egr2*^*+/+*^ *Egr3*^*+/+*^ marrow recipients (**Figure 1H**). However, homozygous *Egr3* deletion combined with heterozygous, and to a greater extent homozygous, *Egr2* deletion was sufficient for a dramatic, cell-autonomous increase in CD21^low^ CD23^low^ B cells (**Figure 1H**). Together, these data demonstrate that *Egr2* and *Egr3* act redundantly, in a cell-intrinsic manner, to suppress the accumulation of CD21^low^ CD23^low^ B cells.

### EGR2/3-deficient CD21^low^ CD23^low^ cells share gene expression profiles with CD21^low^ age- and disease-associated B cells

We performed single-cell RNA sequencing analysis on follicular, CD21^low^ CD23^low^, and B1a CD45.1^+^ or CD45.2^+^ B cells sorted by fluorescence-activated cell sorting (FACS) from the mixed chimeras above (**Figure 2A, Supplementary Figure 4A**). Cell type and *Egr2/Egr3* genotype were the main components of gene expression variation between sorted populations (**Figure 2B, Supplementary Figure 4B**). We began by comparing CD21^low^ CD23^low^ B cells to mature follicular B cells, irrespective of *Egr2*/*Egr3* genotypes. The sorted CD21^low^ CD23^low^ B cells differentially expressed many genes previously associated with CD21^low^ atypical B cells in humans and mice (Benedetto et al., 1992, Moir et al., 2001, Moir et al., 2008, Isnardi et al., 2010, Moratto et al., 2006, Warnatz et al., 2002, Hao et al., 2011, Rubtsov et al., 2011, Charles et al., 2011, Terrier et al., 2011, Rakhmanov et al., 2009, Wei et al., 2007, Jenks et al., 2018, Lau et al., 2017, Russell Knode et al., 2017, Rubtsova et al., 2015): increased expression of *Zbtb32, Fcrl5, Sox5, Gas7, Itgam, Cd19, Cd86, Cd80, Zeb2, Lgals1, Fcgr2b, Irf4, Prdm1, Pdcd1* and decreased expression of *Cr2* (CD21), *Fcer2a* (CD23), *Sell* (CD62L), *Icosl, Ighd, Patj, Il4ra, Cd69, Irf8, Bach2, Ets1, Traf5* (**Figure 2C, D**). The CD21^low^ CD23^low^ B cells appeared poised for antibody secretion based on elevated expression of genes that promote plasma cell differentiation (*Zbtb32, Irf4, Zbtb20* and *Prdm1*) and decreased expression of genes that repress plasma cell differentiation (*Bach2, Ets1*). Gene set enrichment analysis (GSEA) comparing the sorted CD21^low^ CD23^low^ and follicular B cells revealed a high skew (family-wise error rate, FWER, p = 0.000-0.013) towards most genes in sets previously defined as differentially expressed in age-associated B cells (B220^+^ CD93^-^ CD43^-^CD21^-^CD23^-^in GSE81650 (Russell Knode et al., 2017); B220^+^ CD19^+^ CD11b^+^ in GSE28887 (Rubtsov et al., 2011)) compared to follicular B cells (**Figure 2E, Supplementary Figure 4C, Supplementary Tables 1&2**). At the leading edge of genes increased in CD21^low^ CD23^low^ B cells were genes increased in both studies of age-associated B cells: *Bhlhe41, Capn2, Sox5, Adora2a, Fah, Rhobtb1, Sirpa, Timp2, Aldh3b1, Adssl1, Gas7, Rbm47, Pon3, Sspn, Zbtb32, Spaca9, Racgap1, Itgb1, Ak8, Cd9, Itgam, Lgals1, Cd300lf, Adgre1, Cdk14*. At the leading edge of genes decreased in CD21^low^ CD23^low^ B cells were genes decreased in both studies of age-associated B cells *Fcer2a, Bmyc, Gpr174, Tnik, Rapgef4, Tmem108, Cr2, Fchsd2, Zfp318, Satb1, Trim59, Crisp3, Pxk, Lrrk2, Dusp4, Sesn1, Neurl3, Mapk11, Sell, Bnip3, Car2, Rasgef1b, Zfp608, Samsn1, Lfng, Pip5k1b*. These data demonstrate that the CD21^low^ CD23^low^ cells sorted from our mixed chimeras resemble CD21^low^/age-associated/atypical memory B cells reported previously in mice and humans. We therefore refer to them hereafter as CD21^low^ B cells.

**Figure 2.**
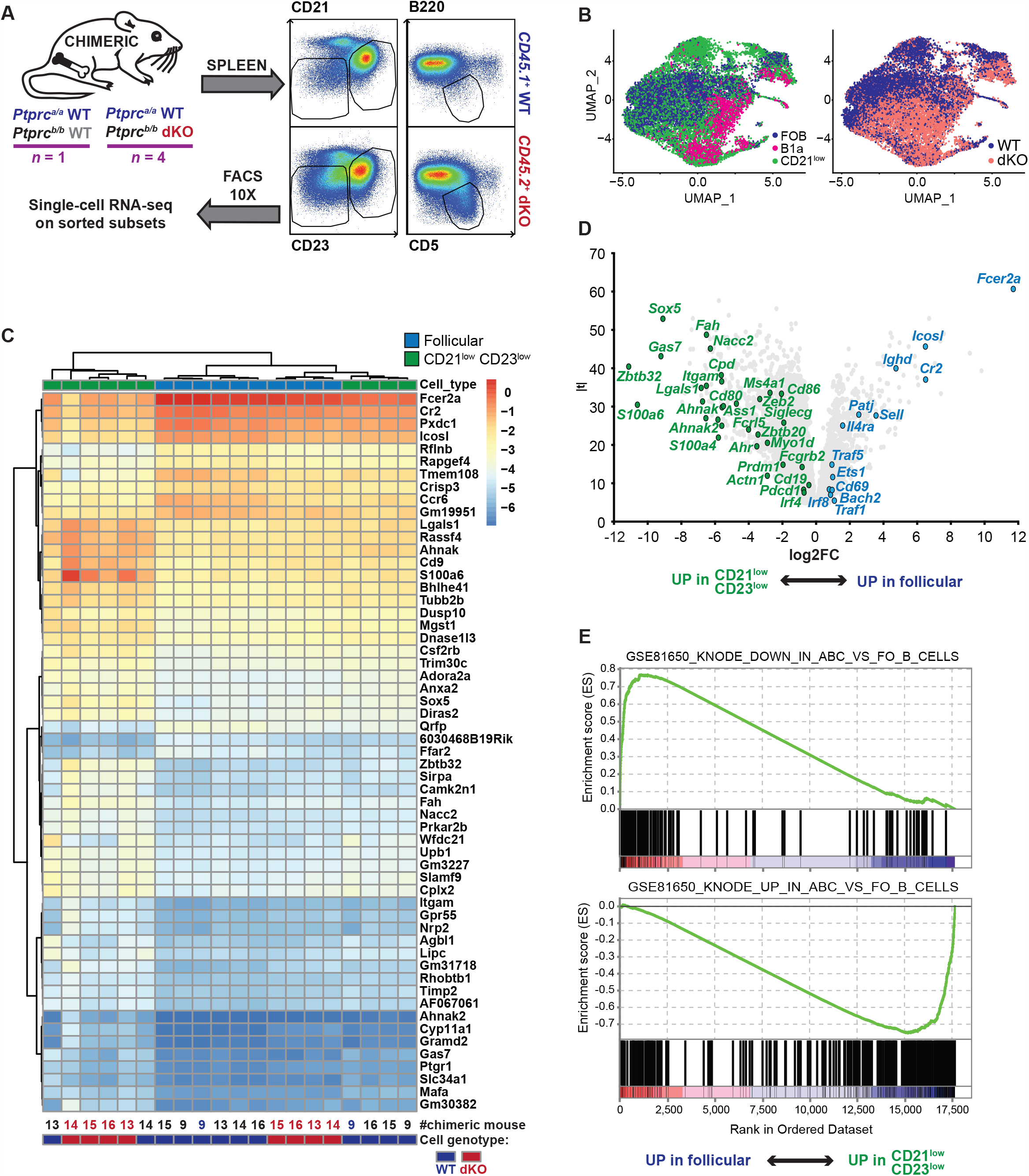
Similarities between CD21^low^ CD23^low^ B cells and *bona fide* age-associated CD21^low^ B cells. **A**, Schematic workflow for single-cell RNA sequencing analysis of sorted CD45.1/CD45.2^+^ splenic B1a (CD19^+^ B220^low^ CD5^+^), CD95^-^CD93^-^mature follicular (CD23^+^) and CD21^low^ CD23^low^ B cells from mixed chimeras transplanted with bone marrow from a *Ptprc*^*a/a*^ *Egr2*^*+/+*^ *Egr3*^*+/+*^ (WT) donor and a *Ptprc*^*b/b*^ WT or *Egr2*^*fl/fl*^ *Egr3*^*KO/KO*^ (dKO) donor. **B**, Unsupervised Uniform Manifold Approximation and Projection (UMAP), following dimensionality reduction of the single-cell RNA short-read sequencing. **C**, Heat map displaying all differentially expressed genes with |log2 fold-change| > 1, between WT (dark blue) and dKO (red) CD21^low^ CD23^low^ (green) and follicular (blue) B cells. **D**, Volcano plot of log2 expression fold-change (log2FC) versus moderated *t*-statistic for differentially expressed genes in CD21^low^ CD23^low^ relative to follicular B cells, irrespective of *Egr2*/*Egr3* genotype, generated using *limma*. **E**, Rank-ordered genes (x axis) and their enrichment scores (y axis) following gene set enrichment analysis (GSEA) of differentially expressed genes in CD21^low^ CD23^low^ relative to follicular B cells, for immunologic terms generated from a published gene set from mouse “age-associated” B cells (Russell Knode et al., 2017).

We next tested for differential mRNA expression between *Egr2*^*fl/fl*^ *Egr3*^*KO/KO*^ (dKO) and wild-type (WT) sorted CD21^low^ B cells exposed to an identical microenvironment. This revealed the cell-intrinsic effects of *Egr2*/*Egr3* deficiency in CD21^low^ B cells. The transcriptome of dKO cells differed significantly from that of WT cells, such that cell type and *Egr2/Egr3* genotype described the two principal components of variation in pseudo-bulk analysis between samples (**Supplementary Figure 4B**). Differentially expressed genes with FWER < 0.05 comprised of 798 increased and 2687 decreased genes, in dKO relative to WT CD21^low^ B cells (**Figure 3A; Supplementary Tables 3&4**).

**Figure 3.**
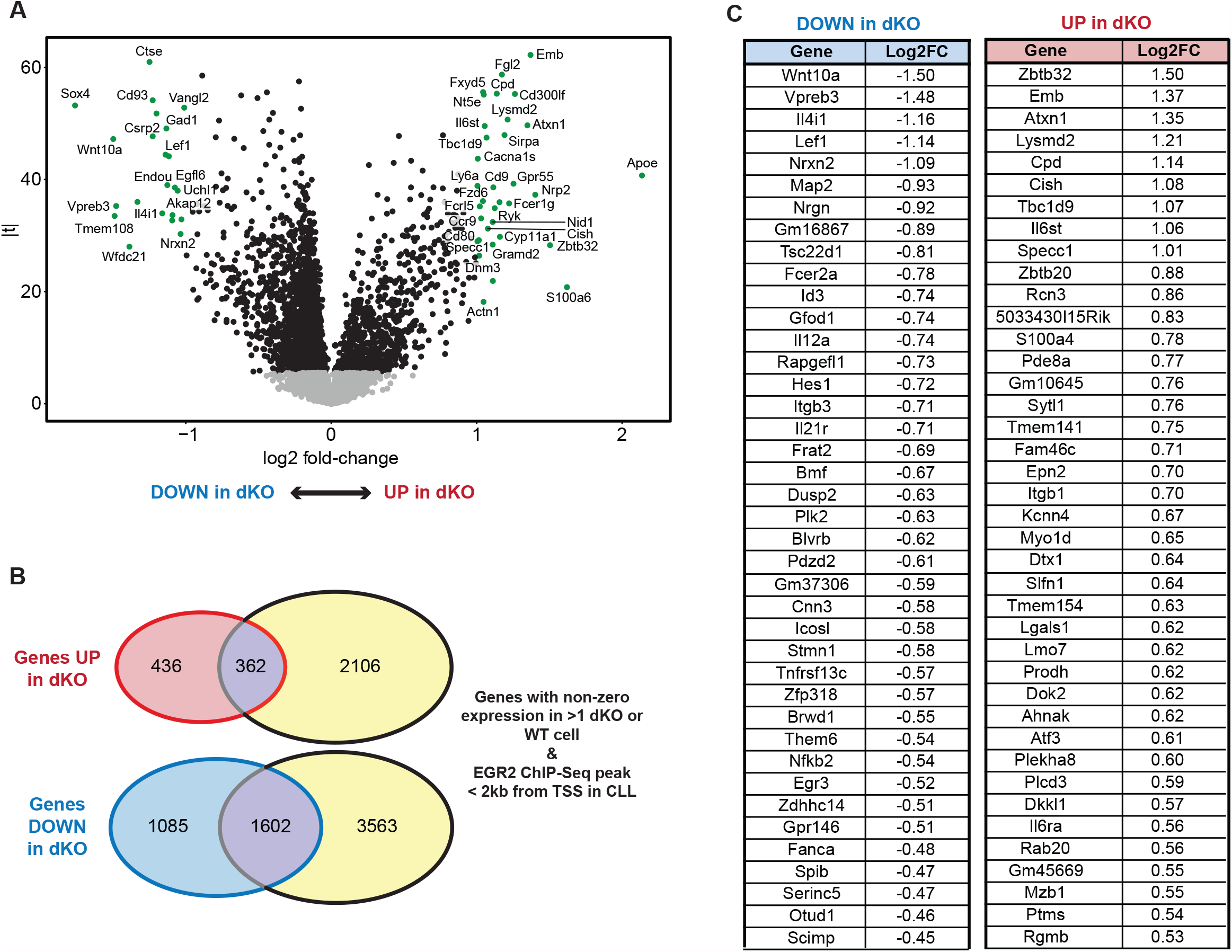
Altered gene expression in *Egr2-* and *Egr3*-deficient relative to wild-type CD21^low^ CD23^low^ B cells, including direct EGR2 targets in CLL. **A**, Volcano plot of log2 expression fold-change (log2FC) versus moderated *t*-statistic for differentially expressed genes in *Egr2*^*fl/fl*^ *Egr3*^*KO/KO*^ (dKO) relative to *Egr2*^*+/+*^ *Egr3*^*+/+*^ (WT) CD21^low^ CD23^low^ B cells, generated using *limma*. Black circles denote genes with a FWER < 0.05. **(B, C)** EGR2 chromatin immunoprecipitation sequencing (ChIP-Seq) was performed independently on *n* = 2 samples of human CLL B cells, and these human hg19 genome coordinates were lifted over to mouse mm10 coordinates. **B**, Venn diagrams displaying genes with non-zero expression in ≥1 cell and increased (red) or decreased (blue) expression in dKO relative to WT CD21^low^ CD23^low^ B cells, that also had ≥1 EGR2 ChIP-seq binding site within 2 kb of their transcription start site (TSS) in ≥1 CLL sample. The enrichment of differentially expressed genes over nearby EGR2 ChIP-seq binding sites was performed using Fisher exact. **C**, Table displaying the top 40 down-regulated (left) or up-regulated (right) genes ranked by |log2FC| in dKO relative to WT CD21^low^ CD23^low^ B cells, that also had ≥1 EGR2 ChIP-Seq binding site within 2 kb of their TSS in human CLL B cells.

Somatic *EGR2* mutations (Damm et al., 2014, Young et al., 2017) and low CD21 expression (Nichols et al., 2015) both correlate with poor prognosis in CLL. To infer which differentially expressed genes in dKO CD21^low^ B cells are direct EGR2 transcriptional targets in B cells, we queried these genes for proximity to EGR2 ChIP-Seq binding domains, with proximity defined as within 2kb from each gene’s transcription start site (TSS), in at least one of two independent human *EGR2-*unmutated CLL samples. On this basis, 362 of the 798 upregulated genes (odds ratio; OR=1.60) and 1602 of the 2687 downregulated genes (OR=1.60) in dKO CD21^low^ B cells have evidence of EGR2 binding in human B cells (**Figure 3B, Supplementary Table 5**). The EGR2 target genes most up-regulated by EGR2/3 deficiency – and hence normally repressed by EGR2/3 - were *Zbtb32, Emb, Atxn1, Lysmd2, Cpd, Cish, Tbc1d9, Il6st, Specc1, Zbtb20* (**Figure 3C**). The EGR2 target genes most down-regulated by EGR2/3 deficiency – and hence normally induced by EGR2/3 - were *Wnt10a, Vpreb3, Il4i1, Lef1, Nrxn2, Map2, Nrgn, Gm16867, Tsc22d1, Fcer2a, Id3*. Many of these direct targets correspond to genes previously identified as differentially expressed by CD21^low^ B cells in mice and humans (Jenks et al., 2018, Rakhmanov et al., 2009, Rubtsov et al., 2011, Russell Knode et al., 2017, Warnatz et al., 2002). Our findings establish EGR2 and EGR3 as major repressors and inducers of gene expression in CD21^low^ B cells.

### EGR2 and EGR3 drive the IgM^low^ IgD^high^ phenotype of self-reactive follicular B cells

Since CD21^low^ B cells in human lupus derive in part from activated follicular B cells (Scharer et al., 2019), we next analysed the expression and role of EGR2 in follicular B cells. Flow cytometric analysis of spleen cells from *Egr2*^*IRES-GFP*^ targeted knock-in reporter mice compared to negative control *Egr2*^*WT*^ mice revealed that the *Egr2* gene is expressed in a large fraction of follicular B cells, correlating inversely with the expression of IgM on their plasma membrane (**Figure 4A**). Cells with highest *Egr2-*GFP expression had the lowest surface IgM whereas there was little detectable *Egr2*-GFP expression in cells with highest surface IgM. The same inverse correlation between *Egr2-*GFP expression and surface IgM existed in immature T1 and transitional T2/3 B cells (**Supplementary Fig 5A**). *Egr2*-GFP expression was higher in B cells from the lowest quartile for cell-surface IgM relative to those from the highest quartile (**Figure 4A, Supplementary Figure 5B, C**). Low cell-surface IgM and high cell surface IgD are due to self-reactivity of the expressed immunoglobulin on mouse and human B cells (Duty et al., 2009, Goodnow et al., 1989, Quach et al., 2011). This finding, that surface IgM down-regulation is a proxy for *Egr2* expression in follicular B cells, extends to single cells in the normal mouse B cell repertoire a previous finding from transgenic mice – that surface IgM is down-regulated and *Egr2* induced as a consequence of chronic binding to self-antigens (Glynne et al., 2000).

**Figure 4.**
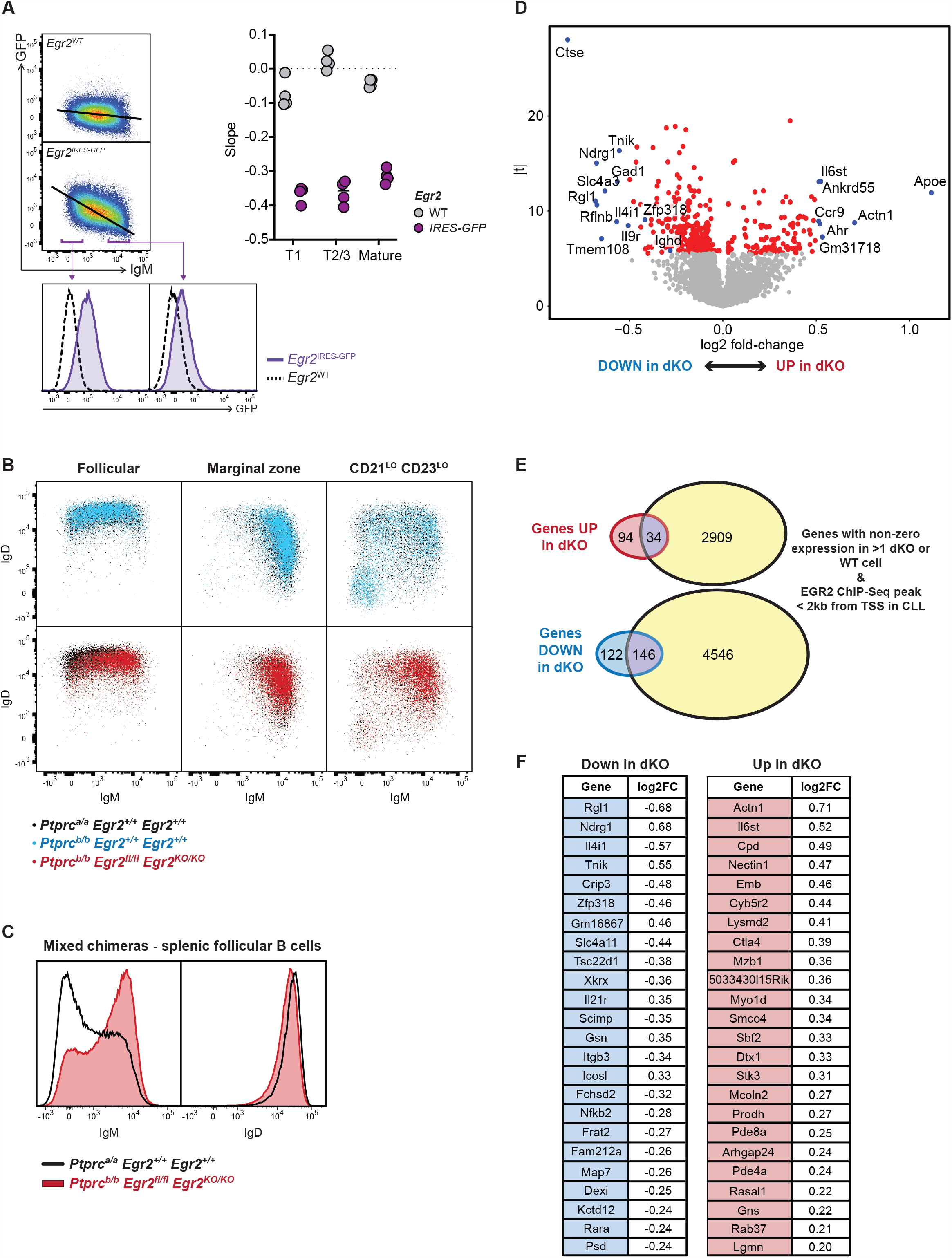
Increased IgM and differential gene expression by *Egr2*- and *Egr3*- deficient follicular B cells. **A**, Representative flow cytometric analysis of IgM versus GFP expression by CD93^neg^ CD23^pos^ follicular B cells from an *Egr2*^*IRES-GFP*^ knock-in mouse and an *Egr2*^*WT*^ reporter-negative control mouse. Right, slope values following linear regression of IgM versus GFP expression in immature T1, T2/3 and follicular B cells from *n*=4 *Egr2*^*IRES-GFP*^ mice (purple) and *n*=4 *Egr2*^*WT*^ reporter-negative mice (grey). Histograms show GFP fluorescence in *Egr2*^*IRES-GFP*^ and *Egr2*^*WT*^ cells within the lowest and highest surface IgM quartiles. **(B-F)** *Rag1*^*KO/KO*^ mice were transplanted with a 1:1 mixture of *Ptprc*^*a/a*^ *Egr2*^*+/+*^ *Egr3*^*+/+*^ (WT, black) and *Ptprc*^*b/b*^ *Egr2*^*+/+*^ *Egr3*^*+/+*^ (WT, blue) or *Egr2*^*fl/fl*^ *Egr3*^*KO/KO*^ (dKO, red) bone marrow. Data representative of *n*=3 experiments with *n*>4 mice per group. **B**, Flow cytometric analysis of IgM versus IgD expression on *Ptprc*^*a/a*^ WT (black) versus *Ptprc*^*b/b*^ WT (blue) or dKO (red) mature B cell populations. **C**, Histogram overlays of IgM and IgD expression on *Ptprc*^*a/a*^ WT (black) versus *Ptprc*^*b/b*^ dKO (red) follicular B cells. **D**, Volcano plot of log2 expression fold-change versus moderated *t*-statistic for differentially expressed genes in dKO relative to WT follicular B cells. Red circles denote genes with a FWER<0.05. **(E**,**F)** EGR2 chromatin immunoprecipitation sequencing (ChIP-Seq) was performed independently on *n*=2 CLL, and human hg19 lifted over to mouse mm10 genome coordinates. **E**, Overlap of genes with increased (red) or decreased (blue) expression in dKO versus WT follicular B cells, that also had ≥1 EGR2 ChIP-seq binding site within 2 kb of their transcription start site (TSS), in ≥1 CLL sample. **F**, Top 24 down-(left) or up-regulated (right) genes, ranked by |log2FC| in dKO versus WT follicular B cells, with ≥1 EGR2 ChIP-Seq binding site <2 kb of their TSS in CLL.

To test for a mechanistic link between *Egr2* expression and low surface IgM, we compared cell-surface IgM and IgD on *Egr2*^*fl/fl*^ *Egr3*^*KO/KO*^ dKO and WT follicular B cells co-existing within mixed chimeric mice. Down-regulation of IgM on most follicular B cells was abolished cell-intrinsically on dKO follicular B cells (**Figure 4B**,**C; Supplementary Figure 6**), accompanied by a more subtle decrease in IgD.

To explore how loss of EGR2/3 causes increased surface IgM, we compared single-cell RNA sequencing data from dKO and WT follicular B cells from mixed chimeras. This revealed 128 increased and 268 decreased genes in dKO follicular B cells with FWER p<0.05 (**Figure 4D, Supplementary Tables 6&7**). Of the 128 increased genes, 34 had EGR2 CHIPseq peaks within 2kb of their TSS in human CLL B cells (**Figure 4E, Supplementary Table 8**, OR=0.81). Of these 34 genes, the 24 genes with the greatest increase in dKO follicular cells (**Figure 4F)** were also all increased in dKO CD21^low^ B cells. Of the 268 genes decreased in dKO follicular B cells, 146 had EGR2 CHIPseq peaks within 2kb of their TSS (**Supplementary Table 8**, OR=1.31). 20 of the 24 most decreased EGR2 target genes in dKO follicular B cells (**Figure 4F)** were also decreased in dKO CD21^low^ B cells, including *Zfp318* (**Supplementary Figure 6B**).

ZFP318 promotes alternative mRNA splicing of the immunoglobulin heavy chain variable (*VDJ*_*H*_) exon to *Ighd* constant region exons instead of the *Ighm* exons, and null or partial loss-of-function *Zfp318* mutations result in aberrantly high surface IgM expression due to increased *Ighm* mRNA and loss of competing IgD protein during assembly with CD79aß (Enders et al., 2014). Consistent with diminished *Zfp318* in dKO follicular B cells, *Ighd* mRNA was also among the decreased gene set (**Supplementary Table 7**).

Collectively, the data above indicate that *Egr2* induction by chronic self-antigen binding in follicular B cells promotes transcription of *Zfp318* to decrease IgM and increase IgD, driving the IgM^low^ IgD^high^ response characteristic of anergic B cells (Duty et al., 2009, Goodnow et al., 1989, Merrell et al., 2006, Quach et al., 2011).

### EGR2 and EGR3 repress B1 cells

B1 B cells exhibit chronic signalling through self-reactive BCRs (Wong et al., 2002, Okamoto et al., 1992, Hayakawa et al., 1999, Graf et al., 2019) and comprise the majority of lymphocytes in the peritoneal cavity, but are infrequent in the circulation and spleen of most mouse strains except the autoimmune prone NZB and NZB/W strains (Hayakawa et al., 1983). Compared to wild-type controls, the mean number of peritoneal CD5^pos^ B1a and CD5^neg^ B1b cells was increased 15- and 16-fold, respectively, in *Egr2*^*fl/fl*^ *Egr3*^*KO/KO*^ mice (**Figure 5A**). A 7-fold increase in mean B1a and B1b cell numbers occurred in *Egr2*^*fl/fl*^ *Egr3*^*KO/+*^ but not in *Egr2*^*fl/fl*^ *Egr3*^*+/+*^ mice (**Figure 5A**). *Egr2*^*fl/fl*^ *Egr3*^*KO/KO*^ mice also had a large increase in mean number of CD19^pos^ B220^int^ CD23^low^ CD43^high^ CD11b^high^ CD5^pos^ B1a cells in the spleen, bone marrow and blood (**Figure 5B**). In mixed chimeras, there was a 30- and 12-fold increase in mean number of *Egr2*^*fl/fl*^ *Egr3*^*KO/KO*^ dKO relative to *Egr2*^*+/+*^ *Egr3*^*+/+*^ WT peritoneal B1a and B1b cells, respectively (**Figure 5C**,**D**). *Egr2*^*fl/fl*^ *Egr3*^*KO/KO*^ B1a cells were also increased 25-fold relative to their wild-type counterparts in the spleen and blood of mixed chimeras (**Figure 5E**).

**Figure 5.**
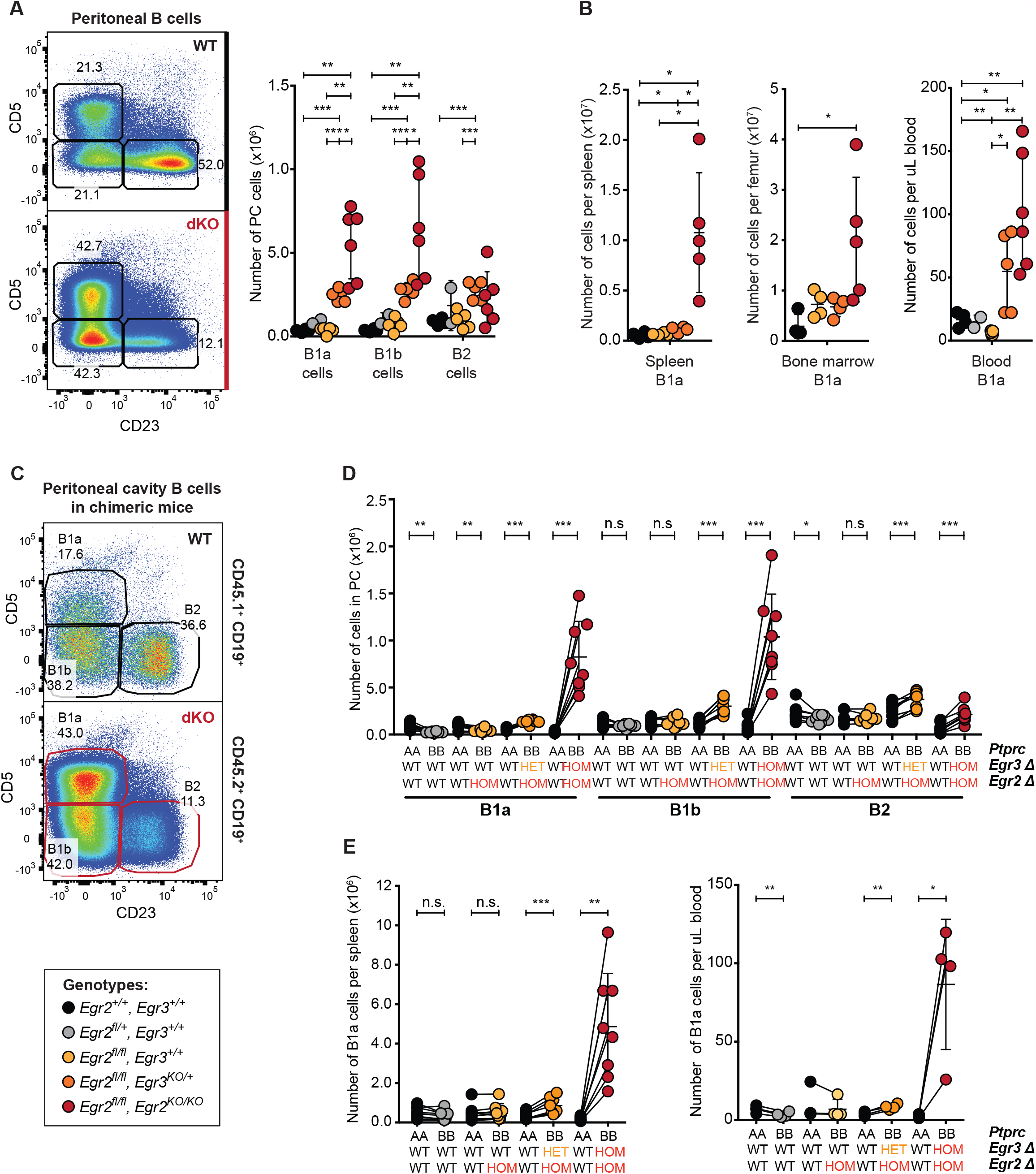
*Egr2-* and *Egr3*-deficiency causes the cell-autonomous accumulation of B1 cells. **A**, Flow cytometric analysis of CD19^pos^ CD23^neg^ CD5^pos^ B1a, CD23^neg^ CD5^neg^ B1b and CD23^pos^ CD5^neg^ B2 cells in *Egr2*^*+/+*^ *Egr3*^*+/+*^ (WT) and *Egr2*^*fl/fl*^ *Egr3*^*KO/KO*^ (dKO) mice (left) and their total numbers in the peritoneal cavity of *Egr2*^*+/+*^ *Egr3*^*+/+*^ (black), *Egr2*^*fl/fl*^ *Egr3*^*+/+*^ (yellow), *Egr2*^*fl/fl*^ *Egr3*^*KO/+*^ (orange) and *Egr2*^*fl/fl*^ *Egr3*^*KO/KO*^ (red) mice (right). **B**, Total number of CD19^pos^ B220^int^ CD23^low^ CD5^pos^ CD43^high^ B1a cells per spleen (left), femur bone (middle) or μL of blood (left), in mice of the indicated genotypes. **(C-E)** *Rag1*^*KO/KO*^ mice were transplanted with a 1:1 mixture of bone marrow from a *Ptprc*^*a/a*^ *Egr2*^*+/+*^ *Egr3*^*+/+*^ donor and from a *Ptprc*^*b/b*^ donor lacking neither, one or both alleles of *Egr2* and/or *Egr3*. Lines link cells within individual chimeric mice. **C**, Flow cytometric analysis of CD45.1^+^ (top) or CD45.2^+^ (bottom) B1a, B1b and B2 cells in the peritoneal cavity of chimeras that received *Ptprc*^*b/b*^ dKO bone marrow. **D**, Total number of *Ptprc*^*a/a*^ or *Ptprc*^*b/b*^ B1a, B1b or B2 cells in the peritoneal cavity of mice that received bone marrow of the indicated genotypes. **E**, Total number of *Ptprc*^*a/a*^ or *Ptprc*^*b/b*^ B1a cells per spleen (left) or per μL of blood (right), in mice that received bone marrow of the indicated genotypes. **(A-E)** Data are represented as mean ± SD. **(A**,**B)** Data representative of *n*=3 experiments in mice 8-20 weeks old. Comparisons made by multiple *t*-tests with Holm-Šidák correction. **(C-E)** Data pooled from *n*=2 independent experiments with *n*=4 recipients per group. Comparisons within a chimeric mouse made by paired t-test. * p < 0.05; ** p < 0.01; *** p < 0.001.

Single cell RNA sequencing of dKO and WT B1a B cells sorted from the spleens of mixed chimeras (**Figure 2A**,**B**) revealed 1667 genes increased and 1783 genes decreased in dKO B1a cells (FWER < 0.05, **Supplementary Tables 9&10**). Strikingly, 354 of 924 genes increased (log2 fold change; log2FC > 0.3) and 257 of 701 genes decreased (log2FC < -0.3) in dKO B1a cells were also differentially expressed using the same threshold in dKO CD21^low^ B cells (**Figure 6A**). Both dKO populations expressed higher levels of genes up-regulated in splenic and peritoneal cavity B1a cells relative to other murine B cell subsets: *Apoe, Ahr, Adm, Cd5, Ctla4, Itgb1, Sox5, Zbtb32*. These populations also had lower expression of genes normally up-regulated upon surface immunoglobulin expression in immature B cells, or as immature B cells develop into follicular B cells, including *Il4il, Zfp318, Xkex, Il21r, Icosl, Fcshd2, Nfkb2*. Notably, EGR2/3-deficient dKO B1a and CD21^low^ B cells overexpressed a set of EGR2-ChIP-seq target genes that are normally increased in cells poised for plasma cell differentiation: *Zbtb32, Zbtb20, Mzb1, Il6st, Il6ra, Pim1, Lgals1*, and *Nfatc1*. More than half of the up-regulated (OR=1.36) or down-regulated (OR=1.60) genes in dKO B1a cells had EGR2 ChIP-seq peaks within 2 kb of their transcription start site in human CLL B cells (**Figure 6B, Supplementary Tables 11**). 253 genes had EGR2 ChIP-seq peaks within 2 kb of their TSS and were increased (log2FC > 0) in both dKO populations, and 617 genes had EGR2 ChIP-seq peaks within 2 kb of their TSS and were decreased (log2FC < 0) in both dKO populations (**Figure 6C**).

**Figure 6.**
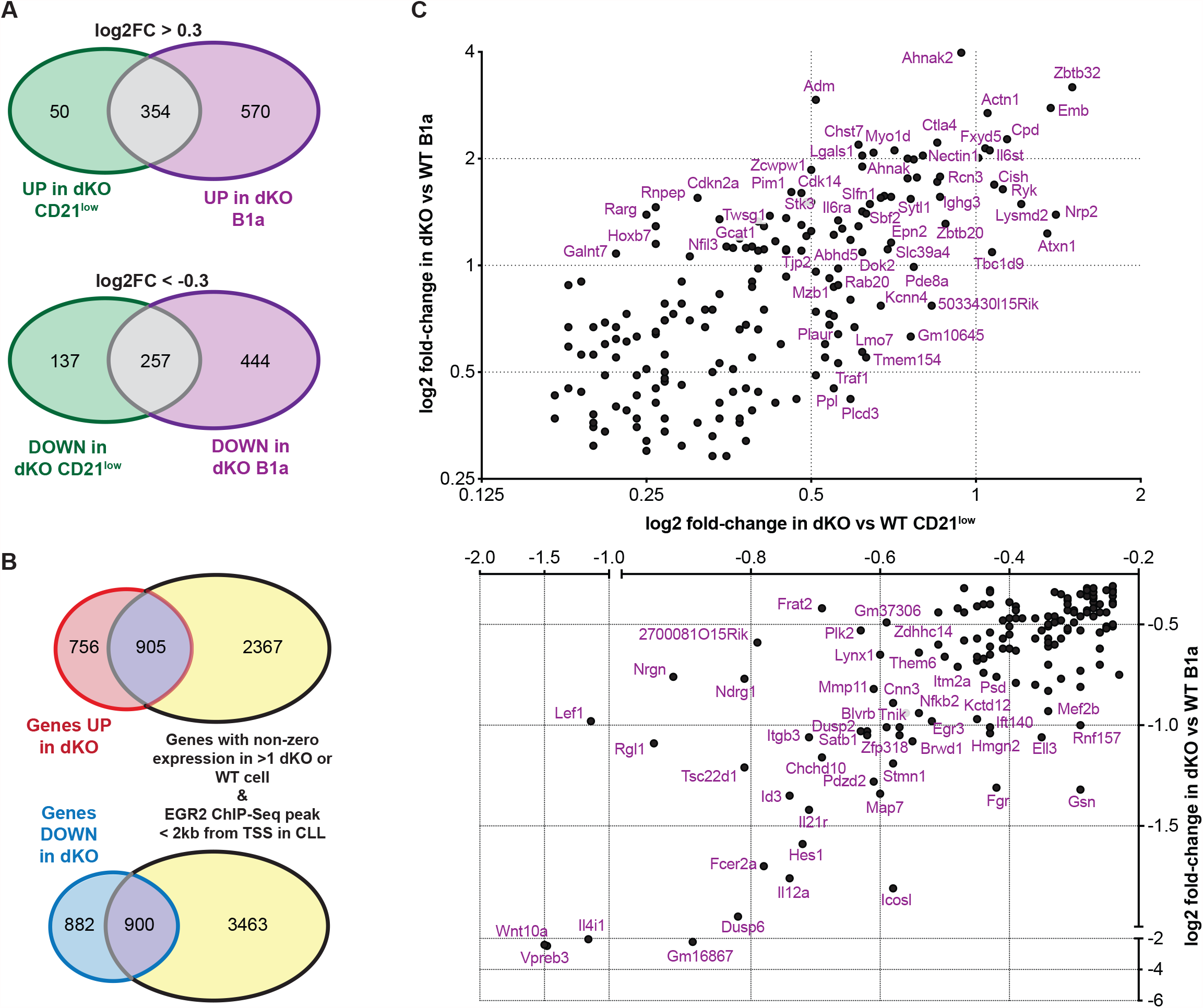
Shared differential gene expression, including of EGR2 targets in CLL, by *Egr2-* and *Egr3*-deficient B1a and CD21^low^ B cells relative to controls. (**A-C**) Single-cell RNA sequence analysis was performed on CD45.1/CD45.2^+^ B1a cells as for other splenic subsets in Figures 2-4. **A**, Overlap of genes significantly (FWER < 0.05) increased (log2FC>0.3) or decreased (log2FC<-0.3) in dKO relative to WT cells, for both CD21^low^ CD23^low^ (green) and B1a (purple) cells. **B**, Overlap of genes significantly (FWER < 0.05) increased (log2FC>0) or decreased (log2FC<0) in dKO relative to WT B1a cells, with genes with non-zero expression and ≥1 EGR2 ChIP-Seq binding site within 2 kb of their transcription start site (TSS) in ≥1 CLL sample. **C**, Scatter plot of genes with EGR2 binding sites within 2 kb of their TSS that were also within the top 200 most differentially expressed genes in dKO versus WT cells, for both B1a and CD21^low^ CD23^low^ B cells.

Collectively, these data demonstrate that EGR2 and EGR3 are critical regulators of B1a cells, and that the landscape of genes regulated by the EGR2/EGR3 checkpoint is similar in B1a and CD21^low^ disease-associated B cells.

## DISCUSSION

Our findings reveal EGR2 and EGR3 as critical regulators of chronically-stimulated B cell populations enriched for self-reactive BCR specificities: IgM^low^ anergic follicular, CD21^low^ and B1a B cells. We reveal that EGR2/EGR3 are essential inducers and repressors of many genes in B cells, particularly essential to repressing a suite of genes that are highly expressed in disease-associated CD21^low^ B cells and drive plasmablast differentiation. These findings and their implications for dysregulated accumulation of CD21^low^ and B1 cells in autoimmune disease and CLL are discussed below.

In mature follicular B cells, which account for the majority of “naïve” (yet to be stimulated by antigen) circulating B cells, a GFP reporter inserted into the 3’ untranslated region of *Egr2* revealed that many are not naïve as they have induced the reporter. Signalling by cell-surface IgM in response to self-antigens, varying in strength from cell to cell, is the most likely stimulus because *Egr2*-GFP induction is correlated with down-regulation of surface IgM. Self-reactive follicular B cells actively down-regulate their surface IgM and activate a state of anergy proportionally to the fraction of their surface Ig molecules chronically engaged by self-antigen (Goodnow et al., 1989). Similar conclusions have been reached using a GFP reporter for another immediate early IgM-induced gene, *Nr4a1*/NUR77 (Zikherman et al., 2012), and for the self-antigen induced gene *Sdc1*/CD138 (Sabouri et al., 2016). However, EGR2/3 is unique because it not only marks but crucially mediates the IgM-down-regulation response of anergic B cells.

The results here elucidate three key mechanisms of B cell anergy: down-regulation of surface IgM/increased IgD, promotion of germinal centre formation, and inhibition of plasmablast differentiation. IgM^low^ IgD^high^ anergic B cells exhibit ongoing, self-antigen-induced oscillations of intracellular calcium that activate calcineurin to dephosphorylate NFAT for nuclear translocation, while selectively uncoupling IgM/IgD signalling to NF-κB (Healy et al., 1997). However, it was not known if any of the myriad calcium/NFAT activated genes mediate anergy (Glynne et al., 2000). This question is addressed by finding that EGR2/EGR3 deficient B cells not only fail to acquire the IgM^low^ IgD^high^ anergic trait, but one of their most decreased mRNAs is *Zfp318*, an EGR2-bound gene in CLL that is induced in anergic B cells (Sabouri et al., 2016) and required to promote alternative mRNA splicing of the VDJ_H_ exon to IgD constant region exons, at the expense of IgM exons (Enders et al., 2014). Consistent with diminished but not absent *Zfp318* mRNA, the increased IgM and lower IgD on EGR2/EGR3-deficient follicular B cells mirrors cells with partial loss-of-function *Zfp318* mutations (Enders et al., 2014). The normal shift to IgD on follicular B cells attenuates IgM signalling to chronic stimulation by self-antigens, and promotes accumulation of IgM^low^ follicular B cells and their recruitment into germinal centre reactions if they bind a foreign antigen (Sabouri et al., 2016).

Three other EGR2 target genes decreased by EGR2/3 deficiency in follicular B cells were *Nfkb2, Il21r* and *Icosl*. IgM^low^ B cells lacking *Nfkb2* have cell autonomously diminished accumulation in follicles (Miosge et al., 2002), while B cells lacking either *Il21r* or *Icosl* are cell autonomously disadvantaged in participating in germinal center reactions (Liu et al., 2015, Tangye and Ma, 2020). EGR2 and EGR3 thus induce a suite of genes that augment follicular B cell recruitment into germinal centre reactions, explaining how this pathway is enhanced in anergic B cells (Sabouri et al., 2016). Promoting germinal centre hypermutation provides a path to remove self-reactivity from immunoglobulins on follicular B cells, by clonal redemption (Burnett et al., 2019).

The third cardinal feature of anergic B cells - their diminished propensity to form plasmablasts (Sabouri et al., 2016), (Goodnow et al., 1989)– is illuminated by finding that EGR2/3-deficient B cells over-expressed a set of EGR2-ChIP-seq target genes normally increased in cells poised for plasma cell differentiation. *Zbtb32* encodes a DNA-binding protein that cooperates with *Prdm1*/BLIMP1 to silence CIITA and is required for long-lived plasma cells (Yoon et al., 2012); *Zbtb20* is an IRF4-induced gene encoding a DNA binding protein that cooperates with *Prdm1*/BLIMP1 to promote plasma cell differentiation and is required for long-lived plasma cells (Chevrier et al., 2014); *Mzb1* is a BLIMP1- and IRF4-induced gene required for plasma cell accumulation (Andreani et al., 2018). *Il6st* encodes the gp130 signalling subunit of the IL-6 receptor that is required for memory B cells in humans (Schwerd et al., 2017). *Il6ra* encodes the other chain of the IL-6 receptor and *Pim1* encodes an IL-6/STAT3-induced protein kinase required for B cell survival and proliferation (Shirogane et al., 1999). *Lgals1* encodes Galectin-1, which is required for B cell proliferation and plasmablast survival (Tsai et al., 2008). Thus, EGR2 and EGR3 normally repress a suite of key driver genes for pre-plasma cells. They do so not only in follicular B cells, but also in CD21^low^ age-associated and B1 B cells.

The transcriptome of EGR2/3-deficient CD21^low^ CD23^low^ B cells closely resembled that of previously described CD21^low^ B cells. A key question arising from our results is the extent to which deficits in the surface IgM-EGR2/EGR3 pathway contribute to accumulation of CD21^low^ CD23^low^ B cells in human lupus and other autoimmune diseases, or in NZB/W mice. In lupus, these derive in part from activated follicular B cells and contribute to the plasmablast population (Scharer et al., 2019). Inherited mutations or polymorphisms or acquired mutations in *EGR2* itself would likely need to function as dominant negative alleles, given the redundancy demonstrated here between complete null mutations in *Egr2* and *Egr3*. Alternatively, genetic deficits upstream at the level of surface IgM signalling to calcium, calcineurin and NFAT could compromise induction of *EGR2* and *EGR3*.

The dramatic dysregulation of B1 cells in the absence of EGR2 and EGR3 is significant, given that somatic *EGR2* mutations recur in CLL (Damm et al., 2014, Young et al., 2017) and that B1a cells accumulate in autoimmune NZB/W mice (Hayakawa et al., 1983) and mouse models bearing recurrent human CLL mutations (Bichi et al., 2002). Self-reactive transgenic B1 cells can escape clonal deletion by localising to the peritoneal cavity, where they are sequestered from exposure to erythrocyte self-antigens (Okamoto et al., 1992). Mice expressing a transgenic BCR recognising Thy-1 develop Thy-1-specific B1 B cells in the presence – but not in the absence – of self-antigen (Hayakawa et al., 1999). An inducible switch in BCR specificity of mature B2 cells to express a typical B1 cell BCR, that recognises self-antigen phosphatidylcholine, leads them to proliferate and differentiate into B1 cells (Graf et al., 2019).

Similar to anergic follicular B cells, surface IgM signalling in B1 B cells chronically elevates intracellular calcium and activates NFAT (Wong et al., 2002). The finding here of *Nfatc1* overexpression in CD21^low^ and B1a cells lacking EGR2/EGR3 is notable. In the *Tcl1*-driven mouse model of CLL, deletion of *Nfatc1* in B cells leads to loss of *Egr2* expression and dramatically accelerates accumulation of leukemic CD5^+^ B cells (Marklin et al., 2017). Given that NFATC1 is up-regulated in activated and B1 cells (Healy et al., 1997) and required for B1a cell accumulation and CD5 expression (Berland and Wortis, 2003), our results indicate that *Egr2* and *Egr3* induction by NFATC1 acts as a negative feedback mechanism preventing excessive B1a cell accumulation. Somatic *EGR2* point mutations in the zinc finger domain are more frequent in aggressive CLL (Damm et al., 2014, Young et al., 2017), but it remains unclear to what extent these cause a general loss-of-function, dominant-negative activity that blocks compensation by EGR3, or change the transcriptional specificity of EGR2 as a repressor or activator of the genes defined here.

The results here fill a major gap in understanding tolerance checkpoints mediating B cell anergy and preventing CD21^low^ and B1 B cell accumulation, to prevent autoimmunity. The critical role for EGR2 and EGR3 in anergic, CD21^low^, and B1 B cells, and the landscape of genes governed by the calcium-NFAT-EGR2/3 checkpoint defined here, will guide future efforts to treat diseases associated with dysregulated accumulation of these cells.

## Acknowledgements

This work was supported by National Health and Medical Research Council (NHMRC) Program (1113904, to C.C.G.) and Fellowship (1081858, to C.C.G.) grants, the UNSW Cellular Genomics Futures Institute, and by The Bill and Patricia Ritchie Foundation. We thank Dr Jonathan D Powell for generously sharing mouse strains, and the Garvan-Weizmann Centre for Cellular Genomics and the Garvan Biological Testing Facility for providing expert technical services.

## Author contributions

E.M-F., J.H.R. and C.C.G. designed the experiments; E.M-F. performed most of the experiments; J.H.R. performed some early experiments; L.A.M., I.A.P provided mouse models used in this study; T.J.P., C.W. and C.C.O conducted RNA-seq and ChIP-Seq bioinformatics analyses; E.M-F., J.H.R. and C.C.G. interpreted the experiments and wrote the manuscript.

## METHODS

### Mouse handling

All mouse handling and experimental methods were performed in accordance with approved protocols of the Garvan Institute of Medical Research/St Vincent’s Hospital Animal Ethics Committee. All mice were bred and maintained in specific pathogen-free conditions at Australian BioResources (ABR; Moss Vale, Australia) or at the Garvan Institute of Medical Research Biological Testing Facility (BTF). All experiments conformed to the current guidelines from the Australian Code of Practice for the Care and Use of Animals for Scientific Purposes. Mice were genotyped by the Garvan Molecular Genetics (GMG) facility at the Garvan Institute of Medical Research.

### Mouse strains

*Cd19*^*Cre*^ mice (Rickert et al., 1997) were obtained from the Jackson laboratory, Bar Harbor, ME. *Egr2* floxed (*Egr2*^*fl*^) (Taillebourg et al., 2002) and *Egr3* knockout (*Egr3*^*KO*^) mice (Tourtellotte and Milbrandt, 1998) were generously provided by Dr. Jonathan D Powell. They were back-crossed >10 generations onto a C57BL/6Ncrl background and crossed with *Cd19*^*Cre*^ mice.

*Egr2*-IRES-GFP reporter mice (Williams et al., 2017) were used to compare *Egr2* and surface IgM expression. These mice express an internal ribosome entry sequence (IRES) followed by the coding region of green fluorescent protein (GFP) targeted into the 3’ untranslated region of the *Egr2* gene (Williams et al., 2017).

C57BL/6 JAusb (C57BL/6J), C57BL/6 NCrl, B6.JSL-*Ptprc*^*a*^*Pepc*^*b*^ (CD45.1) and B6.129S7-*Rag1*^*tm1Mom*^/J (*Rag1*^*KO/KO*^) mice were purchased from ABR.

### Chimeras

To generate 100% chimeras, age- and sex-matched *Rag1*^*KO/KO*^ mice were irradiated with one dose of 425 Rad from an X-ray source (X-RAD 320 Biological Irradiator, PXI). Recipient mice were then injected with donor bone marrow from *Egr2*^*fl*^ *Egr3*^*KO*^ mice lacking one or both alleles of *Egr2* and/or *Egr3*.

To generate mixed chimeras, age- and sex-matched *Rag1*^*KO/KO*^ mice were irradiated with one dose of 425 Rad and 12 hours later, injected with a 1:1 mixture of *Ptprc*^*a/a*^ bone marrow from B6.JSL-*Ptprc*^*a*^*Pepc*^*b*^ donor mice and of *Ptprc*^*b/b*^ cells from *Egr2*^*fl*^ *Egr3*^*KO*^ donor mice lacking one or both alleles of *Egr2* and/or *Egr3*. The bone marrow cell suspension was depleted of lineage-positive cells (expressing B220, CD3, CD4, CD8, CD11b, CD11c, CD19, LY-6C, LY-6G, NK1.1, TCRβ) prior to injection. Each recipient mouse received 2-6 × 10^6^ donor bone marrow cells injected intravenously.

### Flow cytometry and cell-sorting

Single-cell suspensions were prepared from mouse spleen, bone marrow, inguinal lymph nodes, peritoneal cavity and blood. 1-4 × 10^6^ cells in PBS 2% FCS were transferred into appropriate wells of a 96-well U bottom plate. To prevent non-specific antibody binding, cells were incubated with F_c_ blocking antibody for 20 min at 4°C in the dark. Cells were then incubated with antibodies for 30 min, on ice and in the dark. To fix cells, they were incubated in 10% formalin (Sigma-Aldrich) for 15 min at 4°C, and washed and resuspended in PBS 2% FCS. To stain for intracellular nuclear proteins, cells were fixed and permeabilised using the manufacturer’s instructions and the eBioscience Transcription Factor Staining kit. Stained single-cell suspensions were acquired on the BD LSRFortessa^™^. To determine total numbers of populations in the peritoneal cavity, the totality of harvested cells from the peritoneal cavity wash were acquired on the BD LSRFortessa^™^.

Where appropriate, following extracellular antibody staining, immune populations were sorted by fluorescence-activated cell sorting (FACS) on a FACS Aria III (BD Biosciences).

### Anti-mouse antibodies used for flow cytometric analyses

**Anti-mouse antibodies used for flow cytometric analyses.**
| Cat number | Antibody/dye | Fluorochrome | Company | Clone |
| --- | --- | --- | --- | --- |
| A1310 | 7AAD | N/A | Invitrogen | N/A |
| 17-0051-81 | CD5 | APC | Thermo Fischer | 53-7.3 |
| 557396 | CD11b | FITC | BioLegend | M1/70 |
| 101208 | CD11b | PE | BioLegend | M1/70 |
| 45-0114-82 | CD11c | PerCP/Cy5.5 | eBioscience | N418 |
| 115546 | CD19 | BV510 | BioLegend | 6D5 |
| 115539 | CD19 | BV605 | BioLegend | 6D5 |
| 553818 | CD21/35 | FITC | BD Biosciences | 7G6 |
| 101614 | CD23 | PE/Cy7 | BioLegend | B3B4 |
| 101820 | CD24 | Pacific Blue | BD Biosciences | M1/69 |
| 562768 | CD38 | BV421 | BD Biosciences | Ab90 |
| 553270 | CD43 | FITC | BD Biosciences | S7 |
| 564574 | CD45.1 | BUV737 | BD Biosciences | A20 |
| 110716 |  | APC Cy7 | Biolegend | A20 |
| 110728 |  | PerCP Cy5.5 | Biolegend | A20 |
| 560696 | CD45.2 | PE Cy7 | BD Biosciences | 104 |
| 564616 |  | BUV395 | BD Biosciences | 104 |
| 109824 |  | APC Cy7 | BioLegend | 104 |
| 564449 | CD45R/B220 | BUV737 | BD Biosciences | RA3-6B2 |
| 740877 | CD86 | BV786 | BD Biosciences | GL1 |
| 17-5892-83 | CD93 | APC | eBioscience | AA4.1 |
| 557653 | CD95 | PE Cy7 | BD Biosciences | Jo2 |
| 565988 | IgD | BUV395 | BD Biosciences | 11-26c.2a |
| 559750 | Ig, $\kappa$ light chain | Biotin | BD Biosciences | 187.1 |
| 407308 | Ig, $\lambda$ light chain | PE | BioLegend | RML-42 |
| 406515 | IgM | APC/Cy7 | BioLegend | RMM-1 |
| 405229 | Streptavidin | BV605 | BioLegend | N/A |
| 644814 | T-bet | APC | BioLegend | 4B10 |

### Chromatin Immunoprecipitation sequencing (ChIP-Seq)

ChIP-seq was carried out using 30 μg of chromatin isolated from cryopreserved PBMC samples obtained from two CLL patients and 4 μl of anti-EGR2 antibody (Abcam, cat# ab43020, Lot# GR101477-1). The immunoprecipitated DNA was processed into a standard Illumina ChIP-seq library and sequenced on an Illumina HiSeq instrument (Illumina) generating 48 and 35 million 75-nt single-end (SE75) sequence reads. Chromatin was pooled from both samples to generate the input control sequencing library. Sequence reads were mapped to the genome (hg38) using the BWA algorithm with default settings following standard Illumina purity filtering, duplicate read removal, unique mapping, and allowing ≤2 mismatches, generating 28 and 18 million sequence tags. Normalization was performed by down-sampling the number of tags to the lowest number between samples (18 million). Peak intervals were called using the MACS (Zhang et al., 2008) algorithm with the default cut-off *p* value 10^−7^. Peak filtering was performed by removing false ChIP-seq peaks as defined within the ENCODE blacklist.

### Single-cell RNA sequencing using the 10X platform

To assess the cell-autonomous effects of *Egr2* and *Egr3* deficiency on gene expression in B cells, we sorted CD45.1^+^ *Egr2*^*+/+*^ *Egr3*^*+/+*^ (WT) or CD45.2^+^ *Egr2*^*fl/fl*^ *Egr3*^*KO/KO*^ (dKO) splenic mature follicular, CD21^low^ CD23^low^ and B1a cells from *n* = 4 “test” chimeras. To control for any effects of *Ptprc*^*a/a*^ (CD45.1) versus *Ptprc*^*b/b*^ (CD45.2) expression, we sorted CD45.1^+^ WT and CD45.2^+^ WT cells from *n* = 1 control chimera, and later corrected differential gene expression for any differences observed in this comparison at the modelling stage.

Mouse B cells were bulk sorted from mixed chimeras into Eppendorf tubes containing cold sterile PBS 10% FCS and incubated for 20 min at 4°C with TotalSeq^™^ DNA-barcoded anti-mouse ‘Hashing’ antibodies (BioLegend) at a 1/100 final dilution. The TotalSeq^™^ antibodies contain a mixture of two monoclonal antibodies, both conjugated to the same DNA oligonucleotide, that are specific against mouse CD45 and MHC class I haplotypes – and thus stain all leukocytes from C57BL/6 mice.

During the incubation, cells were transferred into a 96-well round bottom plate, on ice. Following incubation, cells were washed three times in cold PBS 2% FCS and the hashed populations pooled into mixtures for single-cell RNA sequencing using the 10X Genomics platform. The Garvan-Weizmann Centre for Cellular Genomics (GWCCG) performed the 10X capture, and sequencing of resulting cDNA samples, as an in-house commercial service, using the Chromium Single-Cell v2 3’ Kits (10X Genomics). A total of 5,000 to 12,000 cells were captured per reaction.

RNA libraries were sequenced on an Illumina NovaSeq 6000 (NovaSeq Control Software v 1.6.0 / Real Time Analysis v3.4.4) using a NovaSeq S4 230 cycles kit (Illumina, 20447086) as follows: 28bp (Read 1), 91bp (Read 2) and 8bp (Index). HASHing libraries were sequenced on an Illumina NextSeq 500/550 (NextSeq Control Software v 2.2.0.4 / Real Time Analysis 2.4.11) using a NextSeq 60 cycles kit (Illumina, 20456719) as follows: 28bp (Read 1), 24bp (Read 2) and 8bp (Index). Sequencing generated raw data files in binary base call (BCL) format. These files were demultiplexed and converted to FASTQ using Illumina Conversion Software (bcl2fastq v2.19.0.316). Alignment, filtering, barcode counting and UMI counting were performed using the Cell Ranger Single Cell Software v3.1.0 (10X Genomics). Reads were aligned to the mm10-3.0.0 (release 84) mouse reference genomes. Raw count matrices were exported and filtered using the EmptyDrops package in R (Lun et al., 2019).

### DNA-barcoded anti-mouse Hashing antibodies

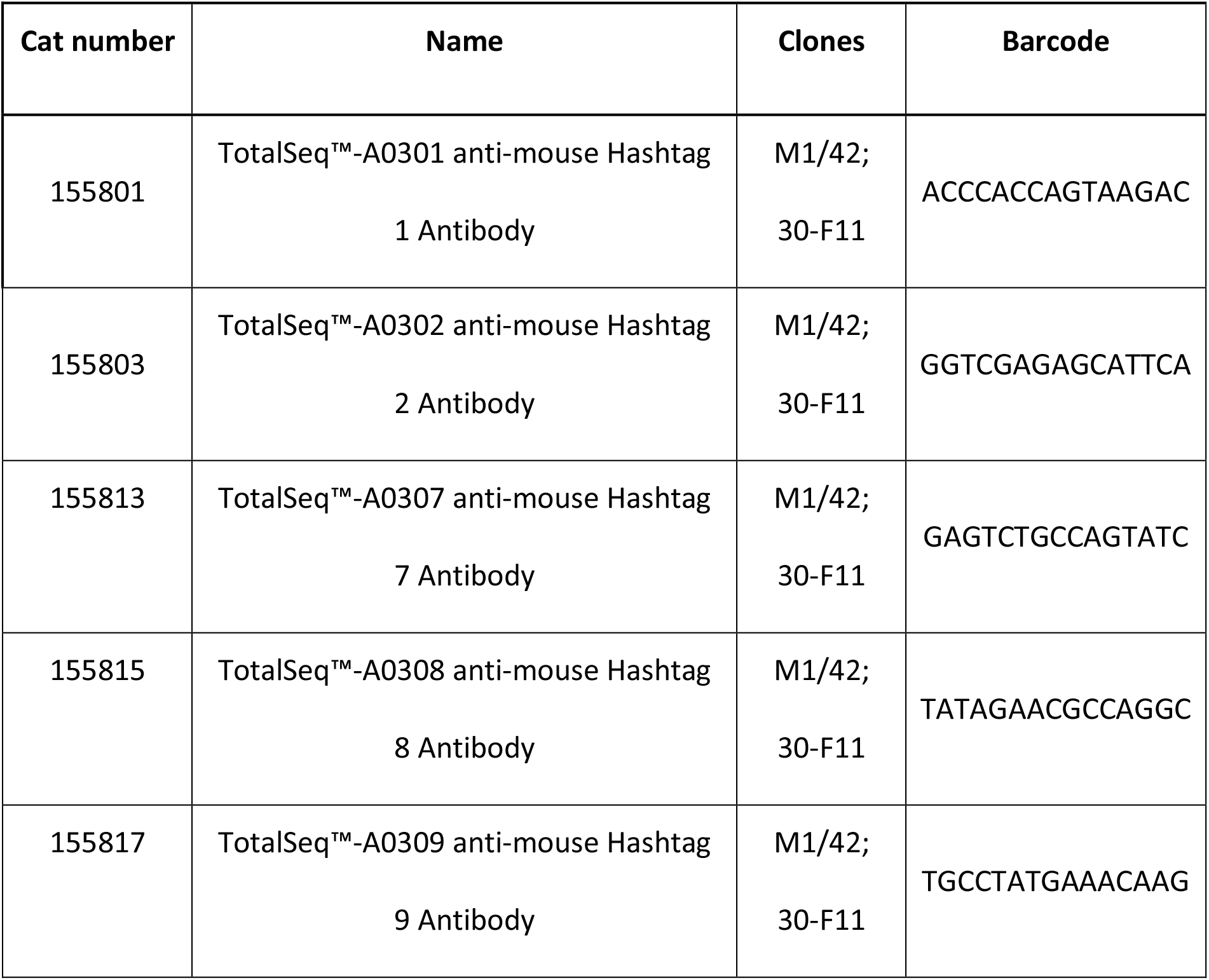
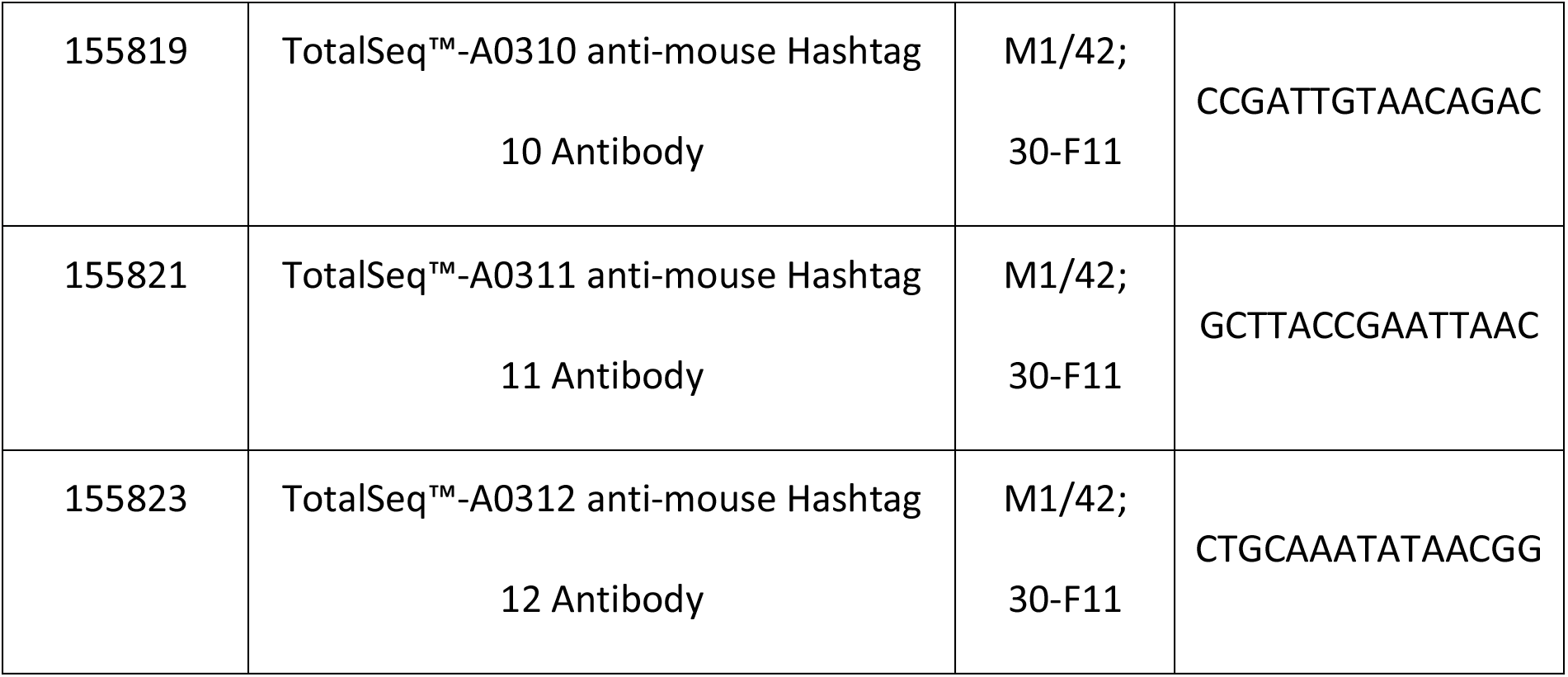

### Statistical analyses

Statistical analyses of flow cytometric experiments were performed using the GraphPad Prism 6 software (GraphPad, San Diego, USA). A one-tailed unpaired Student’s *t*-test with Welch’s correction was used for comparisons between two normally distributed groups. An unpaired student’s *t*-test, corrected for multiple comparisons using the Holm-Sidak method, was used for comparisons of more than two groups. Differences between paired measurements were analysed by paired *t*-test. In all graphs presented, the error bars represent the mean and standard deviation. * p < 0.05, ** p < 0.01, *** p < 0.001.

For the 10X analysis, cells were excluded if the library size or number of expressed genes fell below 2 median absolute deviations, or if mitochondrial reads accounted for more than 20% of total reads. Cell-wise gene expression counts were normalized and recovered using SAVER (Huang et al., 2018) with default values, and differentially expressed genes (DEGs) were identified using limma (Ritchie et al., 2015) on the log-transformed recovered counts. Where appropriate, fold changes and *p*-values were reported after correcting for the *Ptprc* genotype effect during the linear modelling process, through a set of post-hoc contrasts. Bonferroni correction was applied to each set of *p*-values. DEGs were defined as having a family-wise error rate (FWER) < 0.05.

For the EGR2 ChIP-seq analysis, the human genome coordinates of EGR2-bound genes were lifted over to mm10 and a distance below 2 kb from the transcription start site (TSS) was used to define EGR2 peaks in proximity to a given gene. Significant enrichment of differentially expressed (FWER < 0.05) EGR2 target genes identified in at least one independent ChIP-Seq experiment was calculated using a Fisher exact test.

## SUPPLEMENTAL INFORMATION TITLES AND LEGENDS

**Supplementary Figure 1 – No changes in bone marrow B cell populations but increased numbers of splenic immature T1 B cells in *Egr2*- and *Egr3*-deficient mice**.

**A**, Representative flow cytometric analysis of B220^pos^ B cells, IgM^neg^ IgD^neg^ precursor (CD43^pos^ CD24^neg^ pre-pro-, CD43^int^ CD24^int^ pro- and CD43^low^ CD24^pos^ pre-B), IgM^pos^ IgD^int^ immature (CD23^neg^ T1 and CD23^pos^ T2) and IgM^low^ IgD^high^ mature recirculating B cells in the bone marrow of *Egr2*^*+/+*^ *Egr3*^*+/+*^ (top) or *Egr2*^*fl/fl*^ *Egr3*^*KO/KO*^ (bottom) mice. **B**, Percentage of parent population (top) or total number per femur (bottom) of bone marrow B cell populations in *Egr2*^*+/+*^ *Egr3*^*+/+*^ (black), *Egr2*^*fl/fl*^ *Egr3*^*+/+*^ (yellow), *Egr2*^*fl/fl*^ *Egr3*^*KO/+*^ (orange) and *Egr2*^*fl/fl*^ *Egr3*^*KO/KO*^ (red) mice. **C**, Percentage of parent population (top) or total number per spleen (bottom) of splenic B220^pos^ B cells (left), CD93^pos^ immature versus CD93^neg^ mature (middle) and immature CD23^neg^ T1, CD23^pos^ IgM^high^ T2 and CD23^pos^ IgM^low^ T3 (right), in mice of the indicated genotypes. **D**, Representative flow cytometric analysis of splenic immature B cell populations, in *Egr2*^*+/+*^ *Egr3*^*+/+*^ (top) or *Egr2*^*fl/fl*^ *Egr3*^*KO/KO*^ (bottom) mice. (**B**,**C**) Data are represented as mean ± SD. Data are representative of *n* = 3 experiments in mice 8-20 weeks old. Comparisons made by multiple *t*-tests with Holm-Šidák correction. * p < 0.05; ** p < 0.01; *** p < 0.001.

**Supplementary Figure 2 – Reduced B cell CD21 and CD23 expression and accumulation of CD21**^**low**^ **CD23**^**low**^ **B cells in the bone marrow of *Egr2*- and *Egr3*-deficient mice**.

**A**, Left, representative flow cytometric analysis of mature recirculating bone marrow B cell populations, based on CD21 and CD23 cell-surface expression. Right, percentage of CD21^low^ CD23^low^ B cells within mature recirculating B cells in the bone marrow of *Egr2*^*+/+*^ *Egr3*^*+/+*^ (black), *Egr2*^*fl/fl*^ *Egr3*^*+/+*^ (yellow), *Egr2*^*fl/fl*^ *Egr3*^*KO/+*^ (orange) and *Egr2*^*fl/fl*^ *Egr3*^*KO/KO*^ (red) mice. **B**, Mean fluorescence intensity (MFI) of cell-surface antibody staining for CD21 (top) or CD23 (bottom) on bone marrow and splenic B cell populations in mice of the indicated genotypes. (**A**,**B**) Data are represented as mean ± SD. Data are representative of *n* = 3 experiments in mice 8-20 weeks old. Comparisons made by multiple *t*-tests with Holm-Šidák correction. * p < 0.05; ** p < 0.01; *** p < 0.001.

**Supplementary Figure 3 – Accumulation of splenic CD21**^**low**^ **CD23**^**low**^ **B cells in mice transplanted with *Egr2***^***fl/fl***^ ***Egr3***^***KO/KO***^ **bone marrow**.

(**A**,**B**) *Rag1*^*KO/KO*^ mice were irradiated and transplanted with bone marrow cells from a *Ptprc*^*b/b*^ donor mouse that was *Egr2*^*fl/+*^ *Egr3*^*+/+*^ (black) or *Egr2*^*fl/+*^ *Egr3*^*KO/+*^ (light orange) or *Egr2*^*fl/fl*^ *Egr3*^*KO/KO*^ (red). **A**, Frequency of CD19^pos^ B cells, CD93^pos^ immature versus CD93^neg^ mature and of CD23^neg^ T1, CD23^pos^ IgM^high^ T2 and CD23^pos^ IgM^low^ T3 immature B cells, as percentage of splenic leukocytes, in mice transplanted with bone marrow cells of the indicated genotypes. **B**, Left, representative flow cytometric analysis of B220^pos^ CD95^neg^ CD93^neg^ mature B cells with a CD23^pos^ follicular, CD23^low^ CD21^pos^ marginal zone or CD21^low^ CD23^low^ B cells phenotype, in *Egr2*^*+/+*^ *Egr3*^*+/+*^ (top) or *Egr2*^*fl/fl*^ *Egr3*^*KO/KO*^ (bottom) mice. Right, percentage of mature B cells or total number per spleen of follicular, marginal zone and CD21^low^ B cells in mice of the indicated genotypes. (**A**,**B**) Data are represented as mean ± SD. Data representative of *n* = 1 experiment, with *n* = 7 mice per group. Comparisons made by multiple *t*-tests with Holm- Šidák correction. * p < 0.05; ** p < 0.01; *** p < 0.001.

**Supplementary Figure 4 – Schematic workflow and principle components analysis of single-cell RNA sequencing of splenic B cell populations from mixed chimeras**.

**A**, *Rag1*^*KO/KO*^ mice were transplanted with a 1:1 mixture of bone marrow from a *Ptprc*^*a/a*^ *Egr2*^*+/+*^ *Egr3*^*+/+*^ (WT) donor and from an *Egr*^*+/+*^ *Egr3*^*+/+*^ (WT) or *Egr2*^*fl/fl*^ *Egr3*^*KO/KO*^ (dKO) *Ptprc*^*b/b*^ donor. Following reconstitution, CD45.1^/^CD45.2^+^ splenic CD19^pos^ B220^int^ CD5^pos^ CD23^low^ B1a cells, CD19^pos^ B220^pos^ CD95^neg^ CD93^neg^ mature CD23^pos^ follicular or CD21^low^ CD23^low^ B cells were bulk-sorted from 1 “control” chimera that received WT *Ptprc*^*b/b*^ bone marrow and from 4 “test” chimeras that received dKO *Ptprc*^*b/b*^ bone marrow. Each purified population was incubated with a uniquely DNA-barcoded TotalSeq^™^ Hashtag antibody and the barcoded populations were pooled for single-cell RNA sequencing using the Chromium 3’ 10X platform. **B**, Unsupervised analysis using principal components analysis (PCA) of ‘pseudobulk’ gene expression levels in DNA-barcoded populations: B1a cells (squares), mature follicular (circles) and CD21^low^ CD23^low^ (triangles) B cells that were *Ptprc*^*a/a*^ WT (dark blue fill) or *Ptprc*^*bb*^ WT (light blue fill) or *Ptprc*^*b/b*^ dKO (red fill). **C**, Rank-ordered genes (x axis) and their enrichment scores (y axis) following gene set enrichment analysis (GSEA) of differentially expressed genes in CD21^low^ CD23^low^ relative to follicular B cells, for immunologic terms generated from a published gene set from mouse “CD11c^+^” B cells (Rubtsov et al., 2011).

**Supplementary Figure 5 – Negative correlation of cell-surface IgM and *Egr2* gene expression in an *Egr2*-IRES-GFP mouse model**.

**A**, Representative gating and linear regression of cell-surface IgM expression on CD23^neg^ CD93^pos^ immature T1 (left) or CD23^pos^ CD93^pos^ T2/T3 (right) B cells versus green fluorescent protein (GFP) fluorescence, in *Egr2*-IRES-GFP reporter mice. **B**, Representative histogram overlays of GFP expression by T1, T2/T3 or CD93^neg^ CD23^pos^ follicular B cells from *Egr2*^*IRES-GFP*^ mice, gated on cells in the lowest quartile (blue) and highest quartile (red) for cell-surface IgM expression. **C**, Representative histogram overlays of GFP expression by T1, T2/T3 or follicular B cells in the highest (top) or lowest (bottom) quartiles for cell-surface IgM expression, from *Egr2*^*IRES-GFP*^ reporter mice (purple) relative to *Egr2*^*WT*^ reporter negative control mice (black). Data representative of *n*=4 mice per group.

**Supplementary Figure 6 – Altered cell-surface IgM and IgD and *Zfp318* gene expression in *Egr2-* and *Egr3*-deficient B cells**.

(**A-C**) *Rag1*^*KO/KO*^ mice were transplanted with a 1:1 mixture of bone marrow from a *Ptprc*^*a/a*^ *Egr2*^*+/+*^ *Egr3*^*+/+*^ donor and from a *Ptprc*^*b/b*^ donor lacking neither, one or both alleles of *Egr2* and/or *Egr3*. **A**, Mean fluorescence intensity of cell-surface IgM (left) and IgD (right) expression following flow cytometric analysis of B220^+^ CD19^+^ CD95^-^CD93^-^CD23^+^ follicular B cells from mixed chimeras that received bone marrow of the indicated genotypes. Data are represented as mean ± SD. Comparisons within individual chimeric mice were made by paired *t*-test. * p < 0.05; ** p < 0.01; *** p < 0.001. **B**, Violin plots showing kernel density estimations of *Zfp318* gene expression, at single-cell resolution, following single-cell RNA sequencing analysis of *Egr2*^*fl/fl*^ *Egr3*^*KO/KO*^ (dKO; red fill) and *Egr2*^*+/+*^ *Egr3*^*+/+*^ (WT; grey) mature follicular and CD21^low^ CD23^low^ B cells from mixed chimeras. **C**, Representative histogram overlays for IgM (top) or IgD (bottom) cell-surface expression on *Ptprc*^*a/a*^ WT (black line) versus *Ptprc*^*b/b*^ dKO (red fill) splenic B cell populations from chimeric mice. Data representative of *n* > 2 independent experiments with *n* > 4 mice per group.

## SUPPLEMENTARY TABLE TITLES

**Supplementary Table 1:** Gene set enrichment analysis in CD21^low^ CD23^low^ relative to follicular B cell differentially expressed genes, for genes up-regulated in B220^+^ CD93^-^CD43^-^CD21^-^CD23^-^relative to follicular B cells (GSE81650).

**Supplementary Table 2:** Gene set enrichment analysis in CD21^low^ CD23^low^ relative to follicular B cell differentially expressed genes, for genes down-regulated in B220^+^ CD93^-^CD43^-^CD21^-^CD23^-^relative to follicular B cells (GSE81650).

**Supplementary Table 3:** Genes up-regulated in *Egr2/3* dKO relative to WT CD21^low^ B cells.

**Supplementary Table 4:** Genes down-regulated in *Egr2/3* dKO relative to WT CD21^low^ B cells.

**Supplementary Table 5:** Genes differentially expressed in *Egr2/3* dKO relative to WT CD21^low^ B cells, with at least one EGR2 ChIP-Seq peak within 2kb from their transcription start site.

**Supplementary Table 6:** Genes up-regulated in *Egr2/3* dKO relative to WT follicular B cells.

**Supplementary Table 7:** Genes down-regulated in *Egr2/3* dKO relative to WT follicular B cells.

**Supplementary Table 8:** Genes differentially expressed in *Egr2/3* dKO relative to WT follicular B cells, with at least one EGR2 ChIP-Seq peak within 2kb from their transcription start site.

**Supplementary Table 9:** Genes up-regulated in *Egr2/3* dKO relative to WT B1a B cells.

**Supplementary Table 10:** Genes down-regulated in *Egr2/3* dKO relative to WT B1a B cells.

**Supplementary Table 11:** Genes differentially expressed in *Egr2/3* dKO relative to WT B1a B cells, with at least one EGR2 ChIP-Seq peak within 2kb from their transcription start site.

