## Supplemental Figures and Legends for "Uncontrolled CD21^low^ age-associated and B1 B cell accumulation caused by failure of an EGR2/3 tolerance checkpoint"

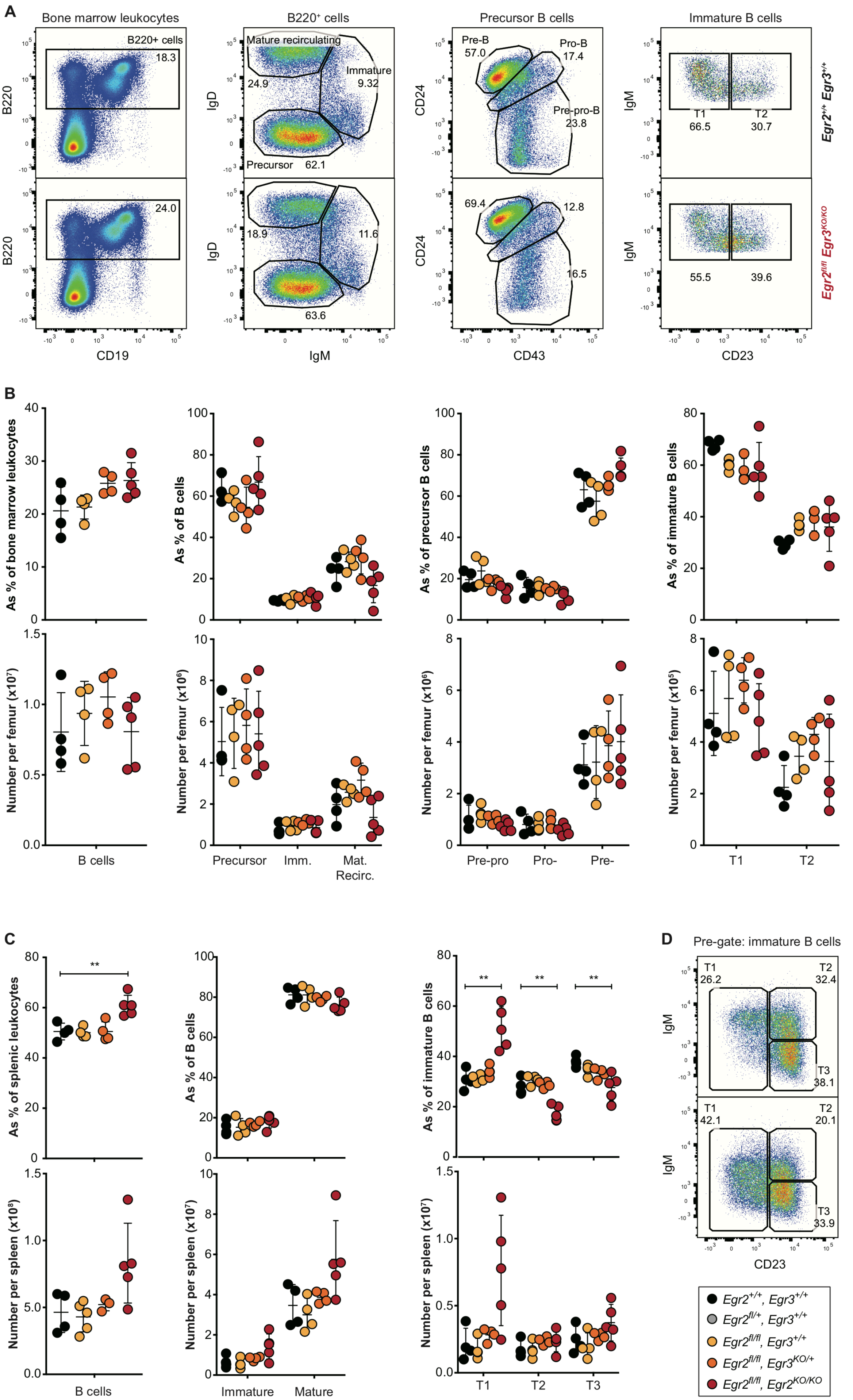
A
Bone marrow leukocytes
B220+ cells
18.3
24.0
B220
CD19
B220⁺ cells
Mature recirculating
24.9
Immature
9.32
Precursor 62.1
18.9
11.6
63.6
IgD
IgM
Precursor B cells
Pre-B
57.0
Pro-B
17.4
Pre-pro-B
23.8
69.4
12.8
16.5
CD24
CD43
Immature B cells
T1
66.5
T2
30.7
55.5
39.6
IgM
CD23
Egr2+/+ Egr3+/+
Egr2fl/fl Egr3KO/KO
B
As % of bone marrow leukocytes
As % of B cells
As % of precursor B cells
As % of immature B cells
Number per femur (x10⁷)
Number per femur (x10⁶)
Number per femur (x10⁵)
B cells
Precursor
Imm.
Mat.
Recirc.
Pre-pro
Pro-
Pre-
T1
T2
C
As % of splenic leukocytes
As % of B cells
As % of immature B cells
Number per spleen (x10⁸)
Number per spleen (x10⁷)
B cells
Immature
Mature
T1
T2
T3
D
Pre-gate: immature B cells
T1
26.2
T2
32.4
T3
38.1
T1
42.1
T2
20.1
T3
33.9
IgM
CD23
Egr2+/+, Egr3+/+
Egr2fl/+, Egr3+/+
Egr2fl/fl, Egr3+/+
Egr2fl/fl, Egr3KO/+
Egr2fl/fl, Egr2KO/KO

**Supplementary Figure 1 – No changes in bone marrow B cell populations but increased numbers of splenic immature T1 B cells in *Egr2*- and *Egr3*-deficient mice.**

**A,** Representative flow cytometric analysis of B220$^{pos}$ B cells, IgM$^{neg}$ IgD$^{neg}$ precursor (CD43$^{pos}$ CD24$^{neg}$ pre-pro-, CD43$^{int}$ CD24$^{int}$ pro- and CD43$^{low}$ CD24$^{pos}$ pre-B), IgM$^{pos}$ IgD$^{int}$ immature (CD23$^{neg}$ T1 and CD23$^{pos}$ T2) and IgM$^{low}$ IgD$^{high}$ mature recirculating B cells in the bone marrow of *Egr2*$^{+/+}$ *Egr3*$^{+/+}$ (top) or *Egr2*$^{fl/fl}$ *Egr3*$^{KO/KO}$ (bottom) mice. **B,** Percentage of parent population (top) or total number per femur (bottom) of bone marrow B cell populations in *Egr2*$^{+/+}$ *Egr3*$^{+/+}$ (black), *Egr2*$^{fl/fl}$ *Egr3*$^{+/+}$ (yellow), *Egr2*$^{fl/fl}$ *Egr3*$^{KO/+}$ (orange) and *Egr2*$^{fl/fl}$ *Egr3*$^{KO/KO}$ (red) mice. **C,** Percentage of parent population (top) or total number per spleen (bottom) of splenic B220$^{pos}$ B cells (left), CD93$^{pos}$ immature versus CD93$^{neg}$ mature (middle) and immature CD23$^{neg}$ T1, CD23$^{pos}$ IgM$^{high}$ T2 and CD23$^{pos}$ IgM$^{low}$ T3 (right), in mice of the indicated genotypes. **D,** Representative flow cytometric analysis of splenic immature B cell populations, in *Egr2*$^{+/+}$ *Egr3*$^{+/+}$ (top) or *Egr2*$^{fl/fl}$ *Egr3*$^{KO/KO}$ (bottom) mice. (**B,C**) Data are represented as mean ± SD. Data are representative of $n = 3$ experiments in mice 8-20 weeks old. Comparisons made by multiple *t*-tests with Holm-Šidák correction. * $p < 0.05$; ** $p < 0.01$; *** $p < 0.001$.

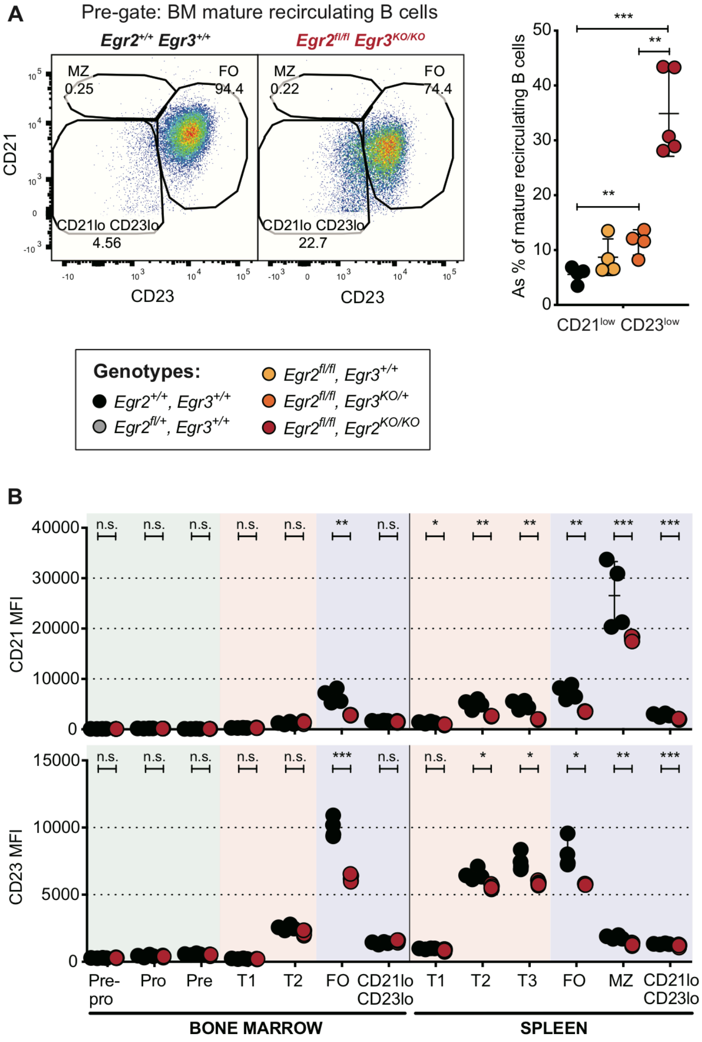


**Supplementary Figure 2 – Reduced B cell CD21 and CD23 expression and accumulation of CD21$^{low}$ CD23$^{low}$ B cells in the bone marrow of *Egr2*- and *Egr3*-deficient mice.**

**A**, Left, representative flow cytometric analysis of mature recirculating bone marrow B cell populations, based on CD21 and CD23 cell-surface expression. Right, percentage of CD21$^{low}$ CD23$^{low}$ B cells within mature recirculating B cells in the bone marrow of *Egr2*$^{+/+}$ *Egr3*$^{+/+}$ (black), *Egr2*$^{fl/fl}$ *Egr3*$^{+/+}$ (yellow), *Egr2*$^{fl/fl}$ *Egr3*$^{KO/+}$ (orange) and *Egr2*$^{fl/fl}$ *Egr3*$^{KO/KO}$ (red) mice. **B**, Mean fluorescence intensity (MFI) of cell-surface antibody staining for CD21 (top) or CD23 (bottom) on bone marrow and splenic B cell populations in mice of the indicated genotypes. (**A**,**B**) Data are represented as mean ± SD. Data are representative of $n = 3$ experiments in mice 8-20 weeks old. Comparisons made by multiple *t*-tests with Holm-Šidák correction. * $p < 0.05$; ** $p < 0.01$; *** $p < 0.001$.

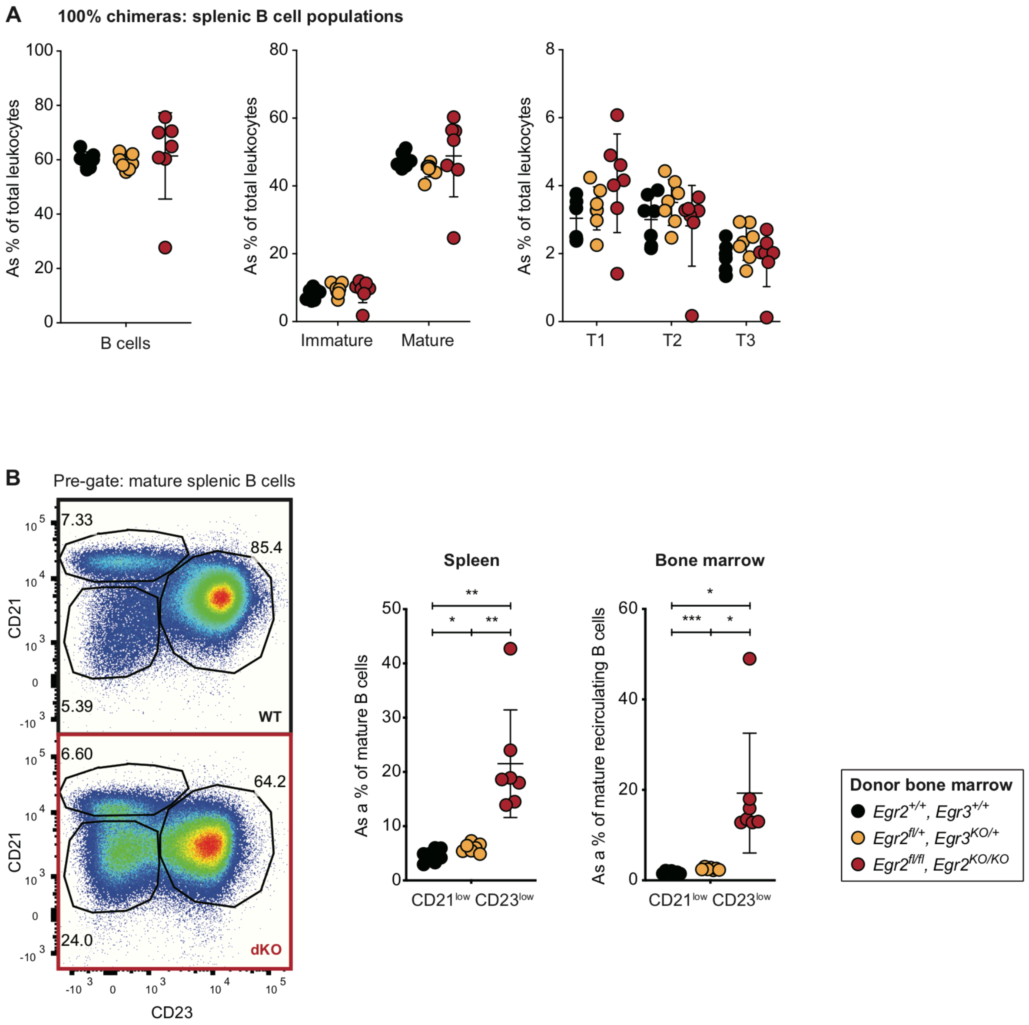


**Supplementary Figure 3 – Accumulation of splenic CD21$^{low}$ CD23$^{low}$ B cells in mice transplanted with *Egr2$^{fl/fl}$ Egr3$^{KO/KO}$* bone marrow.**

(**A,B**) *Rag1$^{KO/KO}$* mice were irradiated and transplanted with bone marrow cells from a *Ptprc$^{b/b}$* donor mouse that was *Egr2$^{fl/+}$ Egr3$^{+/+}$* (black) or *Egr2$^{fl/+}$ Egr3$^{KO/+}$* (light orange) or *Egr2$^{fl/fl}$ Egr3$^{KO/KO}$* (red). **A**, Frequency of CD19$^{pos}$ B cells, CD93$^{pos}$ immature versus CD93$^{neg}$ mature and of CD23$^{neg}$ T1, CD23$^{pos}$ IgM$^{high}$ T2 and CD23$^{pos}$ IgM$^{low}$ T3 immature B cells, as percentage of splenic leukocytes, in mice transplanted with bone marrow cells of the indicated genotypes. **B**, Left, representative flow cytometric analysis of B220$^{pos}$ CD95$^{neg}$ CD93$^{neg}$ mature B cells with a CD23$^{pos}$ follicular, CD23$^{low}$ CD21$^{pos}$ marginal zone or CD21$^{low}$ CD23$^{low}$ B cells phenotype, in *Egr2$^{+/+}$ Egr3$^{+/+}$* (top) or *Egr2$^{fl/fl}$ Egr3$^{KO/KO}$* (bottom) mice. Right, percentage of mature B cells or total number per spleen of follicular, marginal zone and CD21$^{low}$ B cells in mice of the indicated genotypes. (**A,B**) Data are represented as mean ± SD. Data representative of $n$ = 1 experiment, with $n$ = 7 mice per group. Comparisons made by multiple $t$-tests with Holm-Šidák correction. * $p < 0.05$; ** $p < 0.01$; *** $p < 0.001$.

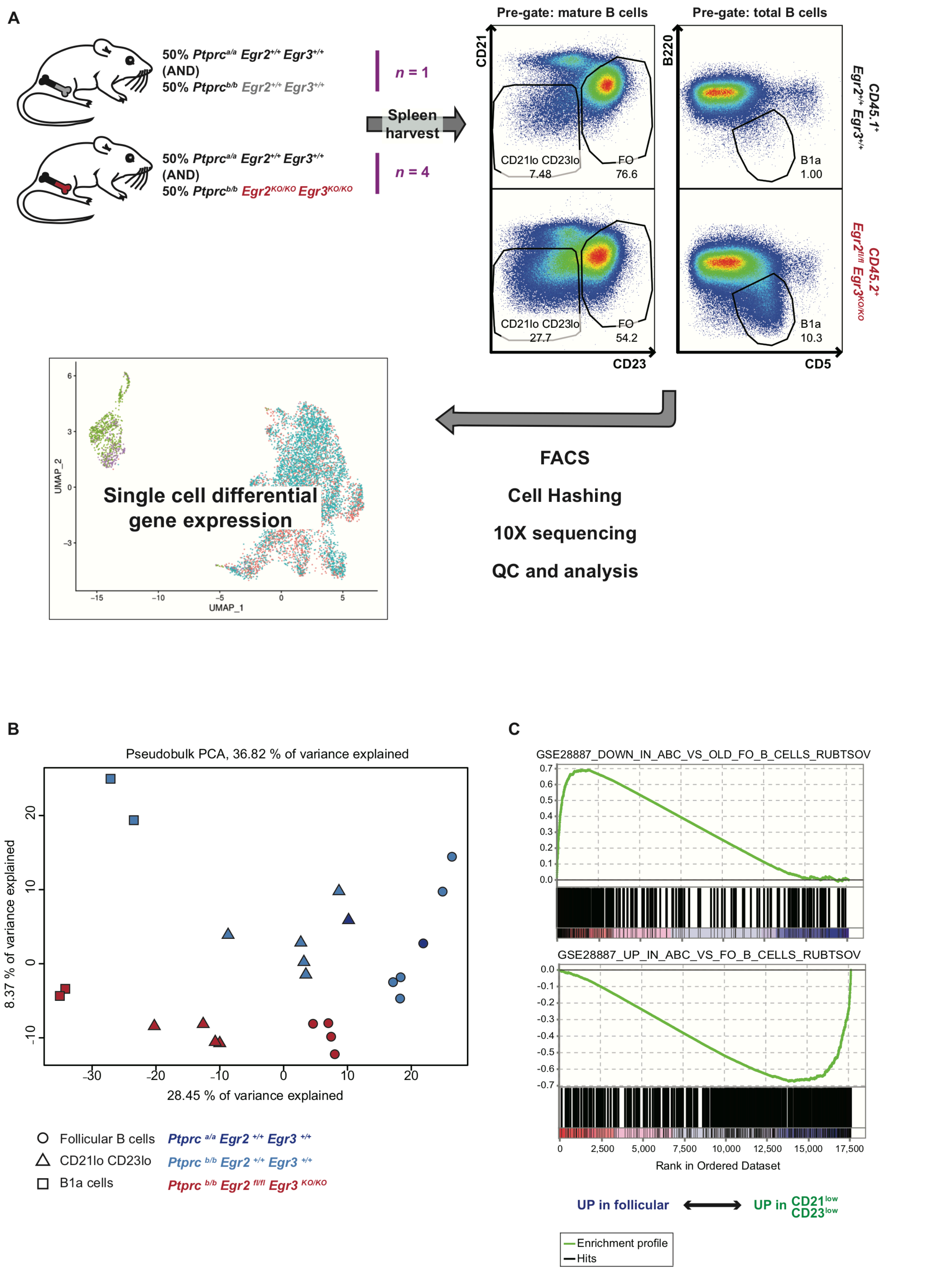


**Supplementary Figure 4 – Schematic workflow and principle components analysis of single-cell RNA sequencing of splenic B cell populations from mixed chimeras.**

**A**, *Rag1$^{KO/KO}$* mice were transplanted with a 1:1 mixture of bone marrow from a *Ptprc$^{a/a}$ Egr2$^{+/+}$ Egr3$^{+/+}$* (WT) donor and from an *Egr$^{+/+}$ Egr3$^{+/+}$* (WT) or *Egr2$^{fl/fl}$ Egr3$^{KO/KO}$* (dKO)

*Ptprc*$^{b/b}$ donor. Following reconstitution, CD45.1/CD45.2$^{+}$ splenic CD19$^{pos}$ B220$^{int}$ CD5$^{pos}$ CD23$^{low}$ B1a cells, CD19$^{pos}$ B220$^{pos}$ CD95$^{neg}$ CD93$^{neg}$ mature CD23$^{pos}$ follicular or CD21$^{low}$ CD23$^{low}$ B cells were bulk-sorted from 1 "control" chimera that received WT *Ptprc*$^{b/b}$ bone marrow and from 4 "test" chimeras that received dKO *Ptprc*$^{b/b}$ bone marrow. Each purified population was incubated with a uniquely DNA-barcoded TotalSeq™ Hashtag antibody and the barcoded populations were pooled for single-cell RNA sequencing using the Chromium 3' 10X platform. **B,** Unsupervised analysis using principal components analysis (PCA) of 'pseudobulk' gene expression levels in DNA-barcoded populations: B1a cells (squares), mature follicular (circles) and CD21$^{low}$ CD23$^{low}$ (triangles) B cells that were *Ptprc*$^{a/a}$ WT (dark blue fill) or *Ptprc*$^{bb}$ WT (light blue fill) or *Ptprc*$^{b/b}$ dKO (red fill). **C,** Rank-ordered genes (x axis) and their enrichment scores (y axis) following gene set enrichment analysis (GSEA) of differentially expressed genes in CD21$^{low}$ CD23$^{low}$ relative to follicular B cells, for immunologic terms generated from a published gene set from mouse "CD11c$^{+}$" B cells [7].

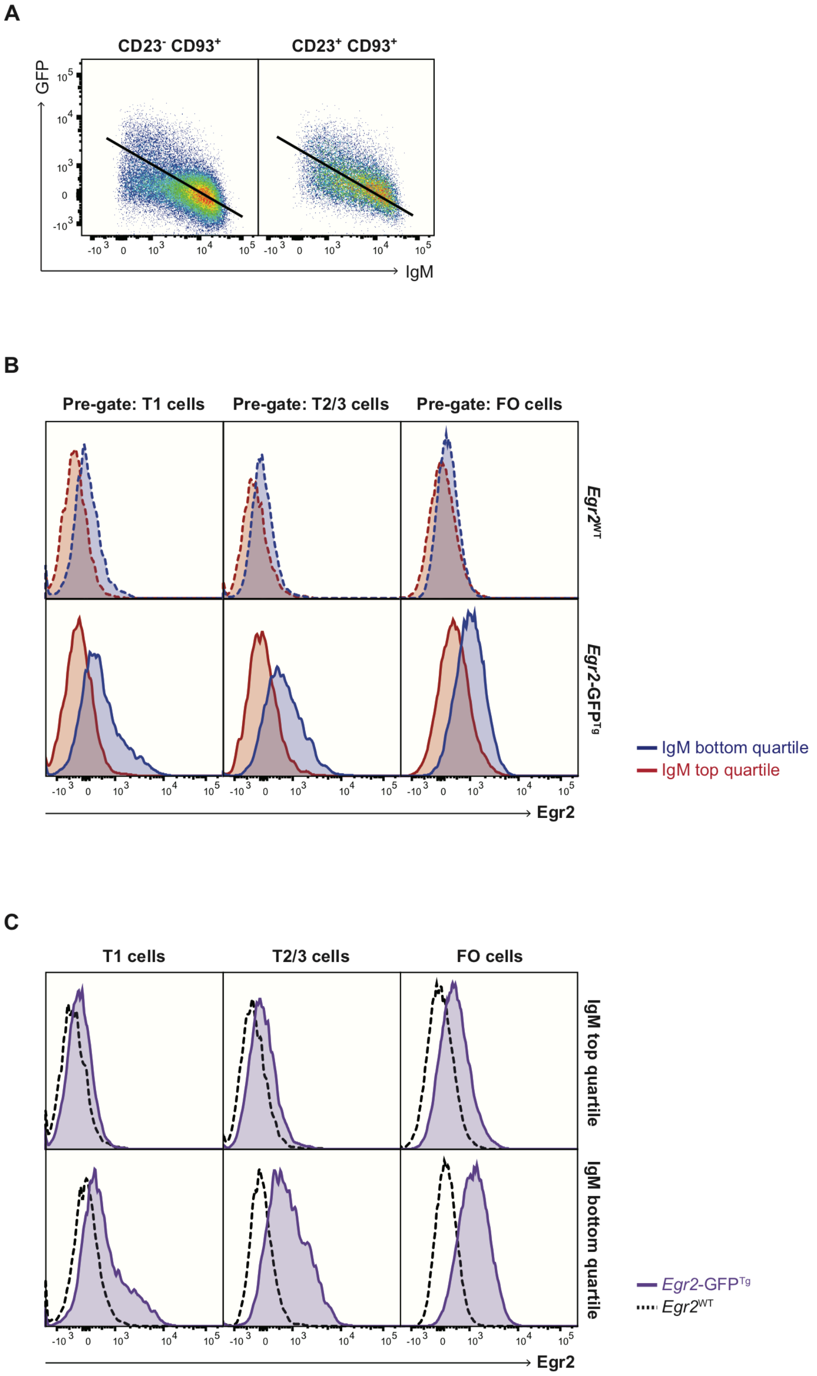


**Supplementary Figure 5 – Negative correlation of cell-surface IgM and *Egr2* gene expression in an *Egr2*-IRES-GFP mouse model.**

**A**, Representative gating and linear regression of cell-surface IgM expression on CD23$^{neg}$ CD93$^{pos}$ immature T1 (left) or CD23$^{pos}$ CD93$^{pos}$ T2/T3 (right) B cells versus green fluorescent protein (GFP) fluorescence, in *Egr2*-IRES-GFP reporter mice. **B**, Representative histogram overlays of GFP expression by T1, T2/T3 or CD93$^{neg}$ CD23$^{pos}$ follicular B cells from *Egr2$^{IRES\text{-}GFP}$* mice, gated on cells in the lowest quartile (blue) and highest quartile (red) for cell-surface IgM expression. **C**, Representative histogram overlays of GFP expression by T1, T2/T3 or follicular

B cells in the highest (top) or lowest (bottom) quartiles for cell-surface IgM expression, from *Egr2^{IRES-GFP}* reporter mice (purple) relative to *Egr2^{WT}* reporter negative control mice (black). Data representative of *n*=4 mice per group.

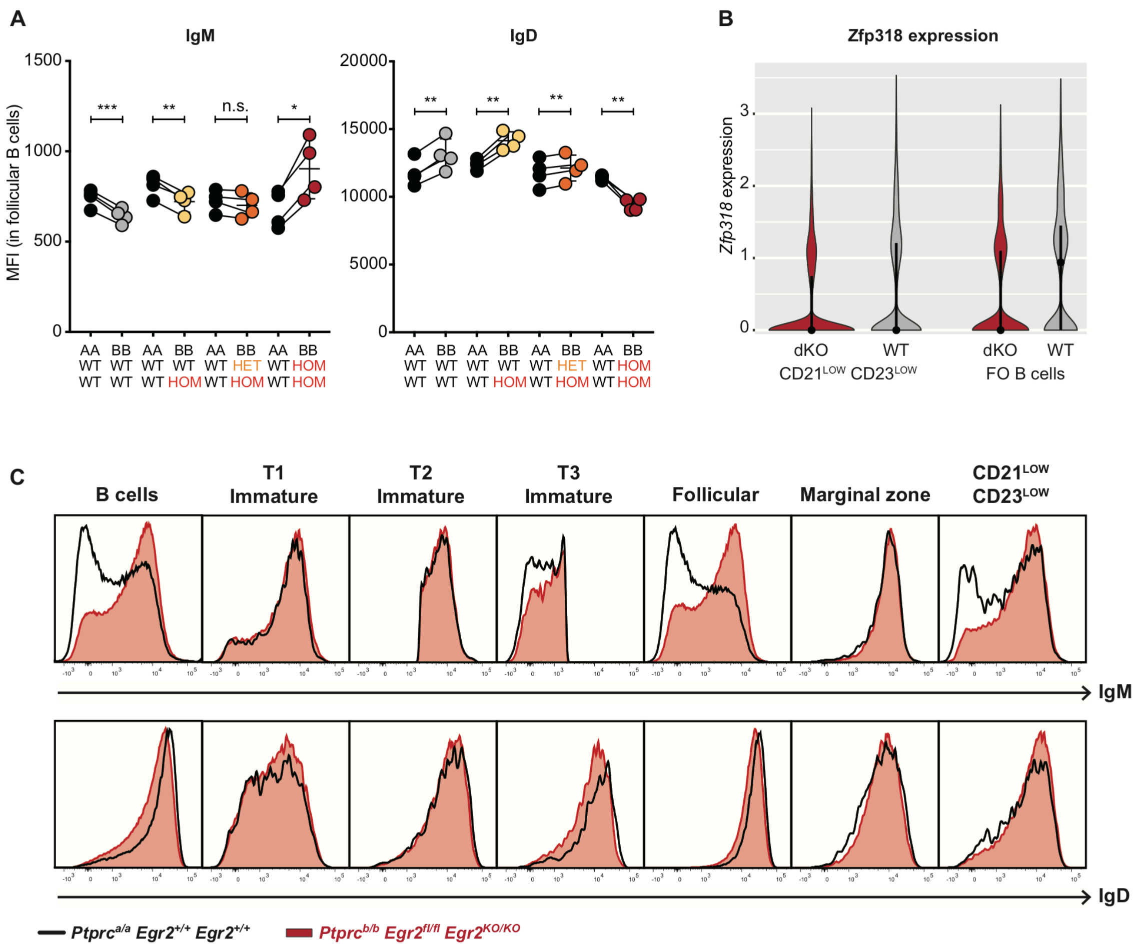


**Supplementary Figure 6 – Altered cell-surface IgM and IgD and *Zfp318* gene expression in *Egr2*- and *Egr3*-deficient B cells.**

(**A-C**) *Rag1^{KO/KO}* mice were transplanted with a 1:1 mixture of bone marrow from a *Ptprc^{a/a}* *Egr2^{+/+}* *Egr3^{+/+}* donor and from a *Ptprc^{b/b}* donor lacking neither, one or both alleles of *Egr2* and/or *Egr3*. **A**, Mean fluorescence intensity of cell-surface IgM (left) and IgD (right) expression following flow cytometric analysis of B220$^+$ CD19$^+$ CD95$^-$ CD93$^-$ CD23$^+$ follicular B cells from mixed chimeras that received bone marrow of the indicated genotypes. Data are represented as mean ± SD. Comparisons within individual chimeric mice were made by paired *t*-test. * $p < 0.05$; ** $p < 0.01$; *** $p < 0.001$. **B**, Violin plots showing kernel density estimations of *Zfp318* gene expression, at single-cell resolution, following single-cell RNA sequencing analysis of *Egr2^{fl/fl}* *Egr3^{KO/KO}* (dKO; red fill) and *Egr2^{+/+}* *Egr3^{+/+}* (WT; grey) mature follicular and CD21$^{low}$ CD23$^{low}$ B cells from mixed chimeras. **C**, Representative histogram overlays for IgM (top) or IgD (bottom) cell-surface expression on *Ptprc^{a/a}* WT (black line) versus *Ptprc^{b/b}* dKO (red fill) splenic B cell populations from chimeric mice. Data representative of *n* > 2 independent experiments with *n* > 4 mice per group.

### SUPPLEMENTARY TABLE TITLES

**Supplementary Table 1:** Gene set enrichment analysis in $CD21^{low}$ $CD23^{low}$ relative to follicular B cell differentially expressed genes, for genes up-regulated in $B220^{+}$ $CD93^{-}$ $CD43^{-}$ $CD21^{-}$ $CD23^{-}$ relative to follicular B cells (GSE81650).

**Supplementary Table 2:** Gene set enrichment analysis in $CD21^{low}$ $CD23^{low}$ relative to follicular B cell differentially expressed genes, for genes down-regulated in $B220^{+}$ $CD93^{-}$ $CD43^{-}$ $CD21^{-}$ $CD23^{-}$ relative to follicular B cells (GSE81650).

**Supplementary Table 3:** Genes up-regulated in *Egr2/3* dKO relative to WT $CD21^{low}$ B cells.

**Supplementary Table 4:** Genes down-regulated in *Egr2/3* dKO relative to WT $CD21^{low}$ B cells.

**Supplementary Table 5:** Genes differentially expressed in *Egr2/3* dKO relative to WT $CD21^{low}$ B cells, with at least one EGR2 ChIP-Seq peak within 2kb from their transcription start site.

**Supplementary Table 6:** Genes up-regulated in *Egr2/3* dKO relative to WT follicular B cells.

**Supplementary Table 7:** Genes down-regulated in *Egr2/3* dKO relative to WT follicular B cells.

**Supplementary Table 8:** Genes differentially expressed in *Egr2/3* dKO relative to WT follicular B cells, with at least one EGR2 ChIP-Seq peak within 2kb from their transcription start site.

**Supplementary Table 9:** Genes up-regulated in *Egr2/3* dKO relative to WT B1a B cells.

**Supplementary Table 10:** Genes down-regulated in *Egr2/3* dKO relative to WT B1a B cells.

**Supplementary Table 11:** Genes differentially expressed in *Egr2/3* dKO relative to WT B1a B cells, with at least one EGR2 ChIP-Seq peak within 2kb from their transcription start site.
